# A novel male accessory gland peptide reduces female post-mating receptivity in the brown planthopper

**DOI:** 10.1101/2024.10.26.620448

**Authors:** Yi-Jie Zhang, Ning Zhang, Ruo-Tong Bu, Dick R. Nässel, Cong-Fen Gao, Shun-Fan Wu

## Abstract

Mating in insects commonly induces a profound change in the physiology and behavior of the female that serves to secure numerous and viable offspring and to ensure paternity for the male by reducing receptivity of the female to further mating attempts. Here, we set out to characterize the post-mating response (PMR) in a pest insect, the brown planthopper (BPH) *Nilaparvata lugens* and to identify a functional analog of sex peptide (SP) and/or other seminal fluid factors that contribute to the PMR in *Drosophila*. We find that BPHs display a distinct PMR that lasts for about 4 days and includes a change in female behavior with decreased receptivity to males and increased oviposition. Extract from male accessory glands (MAG) injected into virgin females triggers a similar PMR, lasting about 24h. Since SP does not exist in BPHs, we screened for candidate mediators by performing a transcriptional and proteomics analysis of MAG extract. We identified a novel 51 amino acid peptide present only in the MAG and not in female BPHs. This peptide, that we designate maccessin (macc), affects the female PMR. Females mated by males with *macc* knockdown display receptivity to wild type males in a second mating, which does not occur in controls. However, oviposition is not affected. Injection of recombinant macc reduces female receptivity, with no effect on oviposition. Thus, macc is so far the only candidate seminal fluid peptide that promotes a PMR in BPHs. Our analysis suggests that the gene encoding the macc precursor is restricted to species closely related to BPHs.

**Author summary:** In insects, mating often induces a long-lasting change in the female behavior and physiology, called a post-mating response (PMR). This ensures numerous and viable offspring, but also serves to secure paternity for the male by inhibiting the female receptivity to further mating attempts. Here, we demonstrate that a pest insect, the brown planthopper (BPH) *Nilaparvata lugens*, also displays a PMR with decreased receptivity to further mating and increased egg laying. We furthermore find that seminal fluid extracted from the male accessory gland of BPHs injected into females generates a PMR. Next, we identified a novel peptide unique to the male accessory gland (designated maccessin) and demonstrate that this peptide is responsible for the reduced receptivity in the PMR, but does not affect egg laying. The gene encoding maccessin appears unique to close relatives of *N. lugens*. This is similar to a *Drosophila* male accessory gland factor, sex peptide, which is known to induce a PMR, and occurs only in a limited number of *Drosophila* species.

## Introduction

Mating in insects commonly leads to a profound change in the physiology and behavior of the female that serves to secure a viable offspring and also to ensure paternity for the male by reducing receptivity of the female to further mating attempts [1–3]. This phenotypic switch has been especially well documented in *Drosophila* where the post-mating response (PMR) includes not only an increase in egg production, but also a reduced receptivity to courting males and changes in feeding, metabolism, and sleep pattern that lasts about a week [1, 3–10]. The trigger of this behavior switch is transferred from the male with the semen during copulation, and in *Drosophila* a major factor is a secreted 36 amino acid peptide, designated sex peptide (SP) [4, 6, 7, 9]. This male-specific peptide, produced in the male accessory gland (MAG) acts primarily on a set of sensory neurons in the female reproductive tract known to express the sex-determination gene *fruitless* and connect to higher order brain circuitry consisting of *doublesex* expressing neurons [11–14]. Thus, transfer of SP and activation of sex-specific neuronal circuits underlie part of the PMR in *Drosophila* females.

Interestingly, SP and the related peptide DUP99B have only been identified in the genomes of a small set of *Drosophila* species and not in other insects [15]. The receptor for SP (SPR) [16] was found to be promiscuous and is additionally activated by myoinhibitory peptide (MIP), also known as allatostatin-B [17, 18]. These authors suggested that MIPs are the ancestral ligands of the SPR, but it is noteworthy that MIPs do not activate the PMR in *Drosophila* [17, 18]. MIPs can be found in most insect species together with its receptor (MIPR). We henceforth use MIPR for this receptor and the *Drosophila* SPR. Although it is possible that MIPs could act as mediators of the PMR in insects that lack SP, there is so far no evidence for this [19].

However, in some insects, it seems that the MIPR is involved in a portion of the PMR as a target of other hitherto unidentified ligands [19]. Examples are the oriental fruitflies *Bactrocera oleae* and *Bactrocera dorsalis* [20–22] and the cotton bollworm *Helicoverpa armigera* [23, 24] where post-mating oviposition is affected by MIPR knockdown. Diminishment of his receptor also affects oviposition in Tobacco cutworm, *Spodoptera litura,* but has no effect on the PMR [25]. The identity of the authentic MIPR ligand(s) remains to be identified in these species.

In other species investigated, the MIPR seems not to be involved in the PMR. The mosquito *Aedes aegypti* is one such case [19]. Interestingly, however, a male-specific peptide was found in the *A. aegypti* MAG and shown to be transferred to females at mating [26]. This decapeptide, *Aedes* head peptide (HP-1), does not act on the MIPR [19, 26, 27]. Instead the HP-1 receptor was identified as a short neuropeptide F receptor (NPYLR1) [28], and interestingly HP-1 induces a life-long refractoriness to insemination by other males [27]. Hence, the mosquito HP-1/NPYLR1 underlies a post-mating change in mate receptivity in females, but has no impact on fecundity, host-seeking or blood-feeding [27], suggesting that this peptide signaling is not fully equivalent to the SP-MIPR axis in *Drosophila*. Finally, there is evidence for a non-peptidergic signal inducing PMR in the malaria mosquito *Anopheles gambiae* [29, 30]. In this species a male-specific form of 20-hydroxyecdysone is sexually transferred to females to induce mating refractoriness [29].

We are interested in the molecular mechanisms and signaling pathway responsible for a possible PMR in a pest insect, the brown planthopper (BPH), *Nilaparvata lugens*. The mating behavior of BPHs has been investigated in some detail [31–34], but it is not yet clear whether females display a post-mating switch in physiology and behavior. Our study identifies a distinct PMR in BPHs with a change in female receptivity to males and an increase in oviposition, lasting for about 4 days. We found that extract from MAGs injected into virgin females induced a similar PMR, although lasting only for about 24h. To screen for a seminal fluid factor responsible for this PMR we performed a transcriptional and proteomics analysis of MAG extract. We identified a novel peptide precursor that turns out to be specific to the MAG in males and not found in female BPHs. The mature 51 amino acid peptide of this precursor was designated maccessin (macc). When exposing females to a second mating after first being mated with males where the *maccessin* (*macc*) gene was knocked down we did not observe any change in receptivity in contrast to controls. However, oviposition is not affected. Injection of recombinant macc peptide reduces female receptivity, but also here there is no effect on oviposition. Thus, we propose that this novel MAG peptide is transferred via seminal fluid to females during copulation and induces a post-mating change in female behavior. Importantly, the gene encoding the macc precursor can only be found in insect species closely related to BPHs. It can be noted that we and a previous study identified another peptide precursor transcript in the MAG [35]. This encodes an isoform (splice variant) of an ion transport peptide (ITPL-1). However, the same ITPL-1 peptide can also be produced by another splice variant in females. We found that knockdown of this peptide in males and injection of recombinant ITPL-1 in females affected the female PMR similar to macc.

## Results

### Brown planthopper females display a distinct post-mating response

Previous studies have described the mating behavior of the brown planthopper (BPH) in some detail [31–34]. During courtship in BPHs the males perform most of the behavioral steps while females only perform a few. The sequence of behaviors includes male abdominal vibration, virgin female abdominal vibration, then males performing following female, wing extension and abdomen vibration, followed by tapping, attempted copulation, copulation and terminated copulation (Figure S1A-H and Video S1). However, there are no reports about a post-mating switch in female behavior and physiology in the BPH. Hence, we first asked whether BPH females display a post mating response (PMR) similar to that observed in *Drosophila* [1, 6, 7, 36, 37], malaria mosquito [29, 30], and other insect species [38–40]. Indeed, we found that once a virgin female BPH has been mated, she is unwilling to accept another courting male (Figure 1A) and lays more eggs than virgin females (Figure 1B). The decreased receptivity of mated females is maintained for at least four days following copulation (Figure 1A). Furthermore, we noted that the PMR in female BPHs includes specific behaviors such as female abdominal vibration and extrusion of the ovipositor towards the courting males, which is also observed in the PMR of *Drosophila* [41, 42] (Figure S1I and J and Video S2). Our data, thus, show that female BPHs exhibit a distinct PMR.

**Fig. 1.**
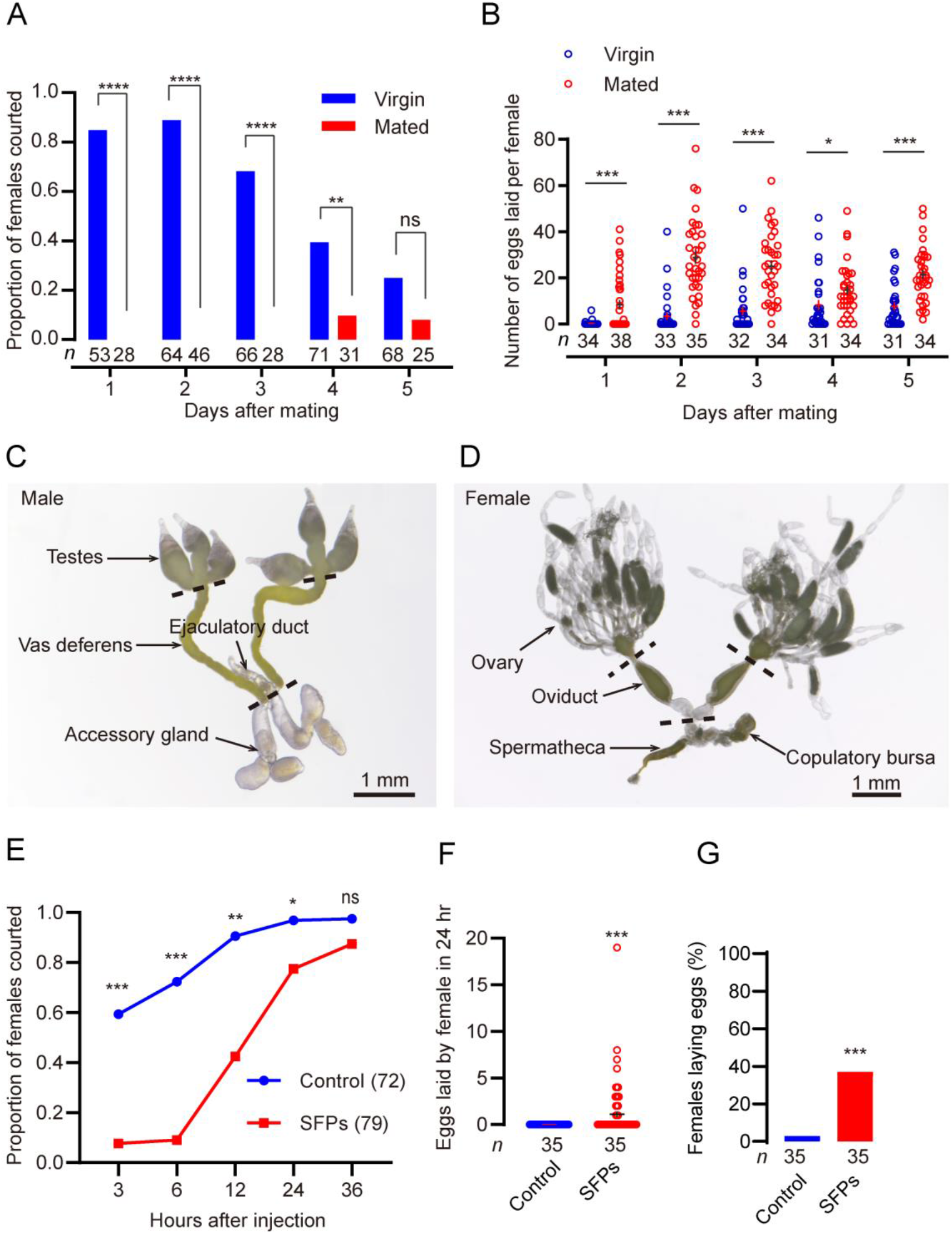
Brown planthoppers display a distinct post mating response and seminal fluid proteins induces a post-mating response. **A.** Female receptivity changes after a first mating. Graph shows the proportion of courted females that respond to males as virgins and mated insects over five days. Rates of courtship differs significantly between mated and virgin females for each of the initial four days. The numbers below the bars denote total number of animals. \*\*\*\**P* < 0.0001, \*\**P* < 0.01, and ns (non-significant), *P* > 0.05; Mann–Whitney test. **B.** Numbers of eggs laid per female. Note that wild type BPH females are known to mate less with age. The small circles and the numbers below the bars denote total number of animals. Data are shown as mean ± s.e.m. Student’s t-test. \**P* < 0.05, \*\**P* < 0.01, \*\*\**P* < 0.001, and ns (non-significant), *P* > 0.05, for comparisons against virgin in A-B. **C**. The reproductive system of the male brown planthopper includes the testes, vas deferens, accessory glands and ejaculatory ducts. **D**. The reproductive system of the female brown planthopper includes the ovary, oviduct, spermatheca and copulatory bursa. **E.** Receptivity to mating in virgin females after injection of male accessory gland proteins extracted from the male BPH, measured as percentage of females that copulated within 30 min. The number of refractory females is high 3-6 h after SFP injection, but then declines. The numbers in brackets denote total number of animals. \**P* < 0.05, \*\**P* < 0.01, \*\*\**P* < 0.001, for comparisons against control; Mann–Whitney test. **F.** Numbers of eggs laid per female in 24 h. The small circles and the numbers below the bars denote total number of animals. \*\*\**P* < 0.001, Student’ t-test. **G.** Percentage of virgins laying eggs during 24 h after SFP injection. The numbers below the bars denote total number of animals. \*\*\**P* < 0.001, Mann–Whitney test (At least four biological replicates with at least five insects per replicate for each experiment).

### Seminal fluid proteins induce a post-mating response in *N. lugens*

Transfer of seminal fluid proteins (SFPs) into virgin females of different insect species, known to display a PMR, results in the repression of female sexual receptivity and stimulates their oviposition to levels similar to those of mated females [2, 5, 7, 16, 37, 43–45]. Hence, we asked whether seminal fluid proteins, transferred into female reproductive organs during copulation could induce a PMR also in BPHs.

Seminal fluid proteins are primarily produced in the male accessary gland (MAG) and ejaculatory duct (Figure 1C) and transferred into female reproductive organs such as copulatory bursa and spermatheca (Figure 1D) during copulation.

Indeed, we found that injection of SFPs, extracted from MAGs of BPHs, into abdomens of virgin females, significantly diminished receptivity to courting males (Figure 1E). These females frequently displayed rejection behavior, e.g. abdominal vibration and ovipositor extrusion, typical of mated females (Video S3). This effect lasts at least 24 hours after injection of SFPs, and thus is shorter than the PMR seen after mating (Figure 1B and 1E). This suggests that there could be other factors that play important roles in long-term PMR in the BPH. Another possibility is that SFPs need to be associated with (bound to) sperm to ensure gradual release of SFPs over a longer duration as was shown in *Drosophila* [46]. We furthermore observed that injection of SFPs into virgin females stimulates egg-laying and leads to an increased percentage (40%) of SFP-injected females ovipositing compared with solvent-injected controls (2%) (Figure 1F and G). Taken together, our results demonstrate that BPHs display a distinct PMR and that SFPs play an important role in this response.

### Transcriptome and proteome analysis of male accessory gland (MAG)

To search for factors that may be transferred with the seminal fluid to induce a PMR, we next performed transcriptional (RNA-seq quantification) and proteomics analyses of MAGs from *N. lugens* (Figure 2-figure supplement 1A).

Illumina sequencing libraries were constructed by using mRNA from the MAG of brown planthoppers. We obtained 54,867,464.7 clean reads on average from samples of MAG (Figure 2-Table S1). After removing low-quality regions, adapters, and possible contamination, we obtained more than 6 giga base clean bases with a Q20 percentage over 98%, Q30 percentage over 94%, and a GC percentage between 38.94 and 41.58% (Table S1). After alignment by Bowtie, 61.01–65.66% and 61.97– 66.21% unique reads were mapped into the reference genome of *N. lugens*. All of the RNA sequence data in this article have been deposited in the China National Center for Bioinformation database and are accessible in CRA019725. To identify the putative function of assembled transcripts, sequence similarity search was conducted against the NCBI non-redundant (NR) and Swiss-Prot protein databases using BLASTx search with a cut-off E value of 10^−5^.

**Figure 2.**
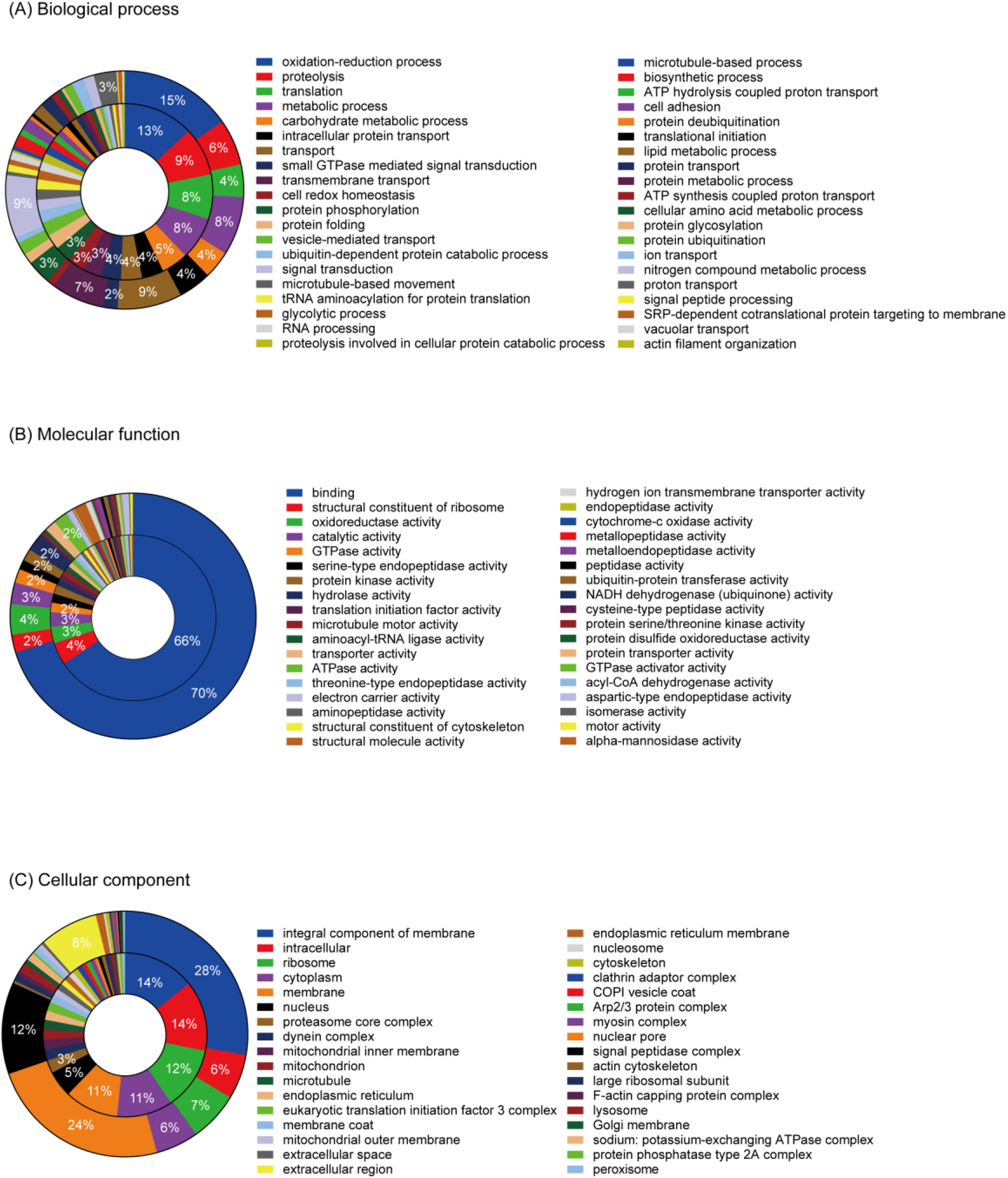
GO classification of brown planthopper MAG transcripts and proteins. Genes and proteins are classified according to gene ontology annotations, and the proportions of each category are shown in terms of percentages of (A) biological processes, (B) molecular functions, and (C) cell components. Outer ring, transcript, inner ring, protein.

Proteomic analysis was performed using Label-free. Production data was searched against brown planthopper MAG transcriptome databases using the Proteome Discoverer 2.2 (PD2.2, Thermo) and identical search parameters. Searching against the de novo assembled *N. lugens* MAG transcriptome, proteomics analysis identified 366,540 total spectra, 101,639 spectra after cleaning and quality checks, 27.73 percent in the total spectra, with the identified 28998 peptides and 3951 proteins. The mass spectrometry proteomics data have been deposited to the Omix of the China National Center for Bioinformation database (https://ngdc.cncb.ac.cn/omix/) via the PRIDE partner repository with the dataset identifier OMIX007634.

Gene ontology (GO), an international standardized gene functional classification system, was used to classify the function of the predicted brown planthopper genes. Based on sequence homology, a total of 16,367 transcripts (36.78%) and 1911 proteins could be categorized into three main categories: biological process, cellular component, and molecular function, with 112 function groups in the transcripts and 112 in the proteins (Figure 2). Genes and proteins involved in oxidation-reduction process was the largest category in biological processes, including 1121 (15.3%) transcripts, 113 (12.9%) proteins. There were 459 (6.3%) transcripts and 78 (8.9%) proteins involved in proteolysis, 72 proteins involved in translation and 69 involved in metabolic process, and the number of proteins involved in signal transduction and transport reached 630 and 644 transcripts, respectively (Figure 2A). In the cellular component category, proteins involved in integral component of membrane (1750, 27.7% transcripts and 79, 14.2% proteins respectively) and membrane (1509, 23.9% transcripts and 59, 10.6% proteins) were all prominently represented (Figure 2C). The genes and proteins associated with ‘binding’ were 70.2% and 65.7% respectively in the molecular function category (Figure 2B). This pattern of distribution is typically seen in the transcriptome of samples undergoing development processes [47]. In our database, 586 transcripts were annotated as related to metabolic processes, which suggests that this analysis provides abundant information on novel genes involved in metabolic pathways, including secondary metabolism.

The Kyoto Encyclopedia of Genes and Genomes (KEGG) database was utilized to categorize gene function and pathways. There were 10,582 transcripts mapped into 229 KEGG pathways. The maps with the highest transcripts representation were signal transduction (1565 transcripts, 14.8%), followed by endocrine system (942 transcripts, 8.9%), carbohydrate metabolism (795 transcripts, 7.5%), and amino acid metabolism (584 transcripts, 5.5%) (Figure 2-figure supplement 2A). The presence of abundant metabolic pathways has also been found in the proteomics of accessory gland of brown planthopper [48]. There were 1047 proteins that mapped into 122 KEGG pathways, with the highest protein representation in global and overview maps (344, 16.5%), followed by carbohydrate metabolism (228, 10.9%), transport and catabolism (202, 9.7%), folding, sorting and degradation (202, 9.7%), translation (195, 9.3%), overview (160, 7.7%), amino acid metabolism (122, 5.8%) (Figure 2-figure supplement 2B).

### Identification of seminal fluid proteins of *N. lugens*

Genes encoding seminal fluid proteins were predicted using both the *N. lugens* transcriptome assembly and proteomic analysis. We identified a total of 373 putative SFPs from the MAG transcriptome data. Of these, 209 sequences were confirmed by proteomic analysis (Figure 2-figure supplement 1B and Table S2). Among of these, 131 putative SFPs have signal peptides and are likely to be secreted by the MAG (Table S2). One of these gene transcripts in the MAG encodes an ion transport peptide-like peptide precursor (ITPL-1) (Table S2). It had previously been reported that one splice isoform of this gene is specifically expressed in the MAG of BPHs [35, 48]. However, the same mature peptide (ITPL-1) could be produced from another splice variant of the same gene in females [35, 48], suggesting that this ITPL-1 signaling can also be endogenous to females.

Interestingly, we discovered a hitherto unknown peptide precursor in the MAG by transcriptome and mass spectrometric analysis (Table S2). This is encoded by the gene BAO00947, which has been previously annotated as a peptide precursor [47].

The peptide encoded on this precursor is 91 amino acids and the mature peptide between the KR-cleavage sites is 51 amino acids long, and has six cysteines that can form three disulfide bridges, spaced in a fashion resembling ion transport peptides [49, 50]. There is no C-terminal amidation signal suggesting that the peptide is non-amidated. We designate this peptide maccessin (<u>m</u>ale <u>access</u>ory gland peptide; <u>macc</u>). In Figure 3A, we show the amino acid sequence of the predicted neuropeptide precursor encoded by BAO00947, including the signal peptide and cleavage sites. A macc peptide fragment (ATLGEYTY) could also be detected in MAG tissue extract in our proteomics analysis (Table S2). We performed a semi-quantitative RT-PCR analysis of *mac*c and found that it is only expressed in the MAG, and cannot be found in males with the MAG removed, or in females (Figure 3B). Thus, *mac*c is a male and tissue-specific peptide.

**Figure 3.**
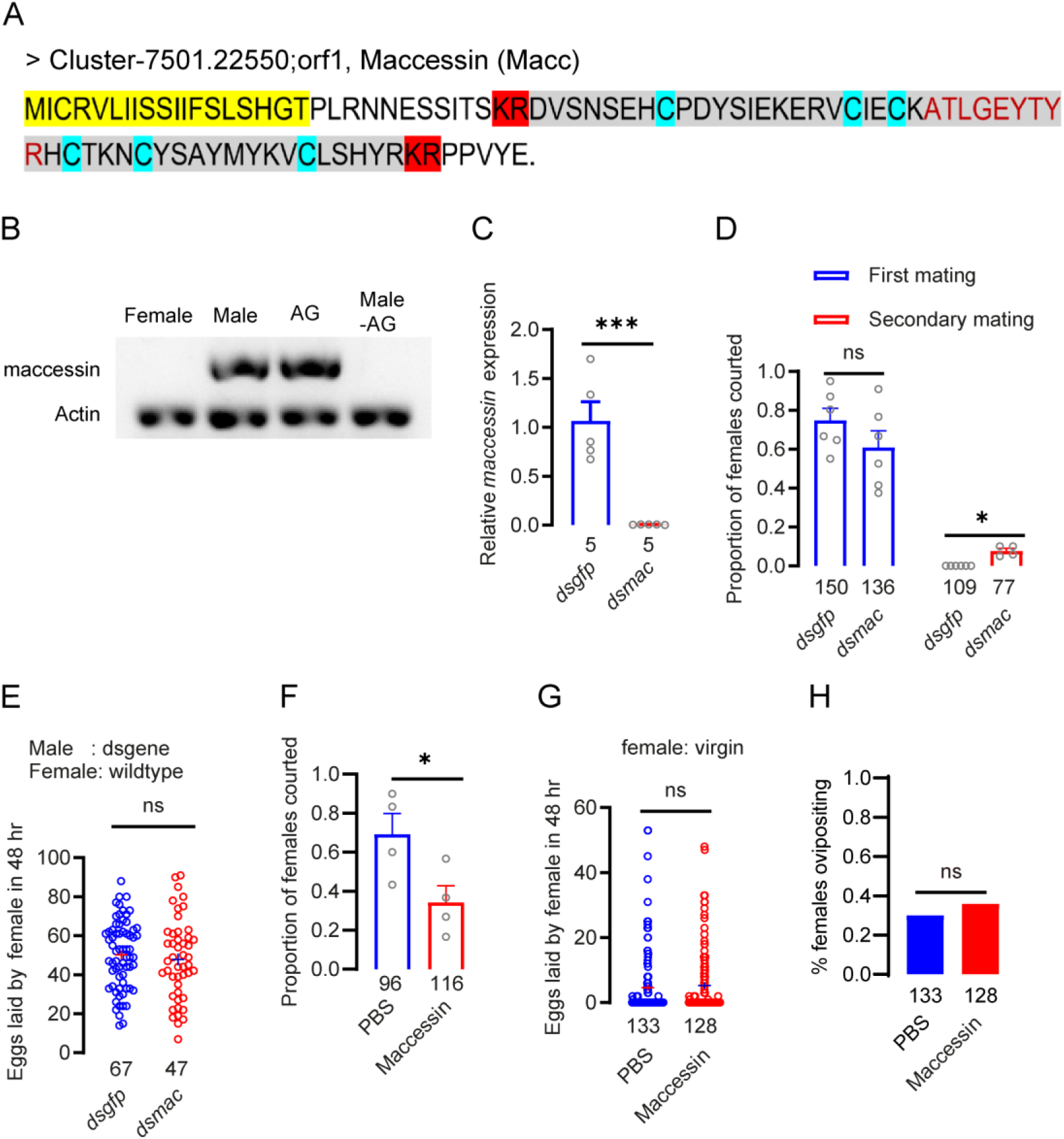
Maccessin reduces female receptivity but not ovipostion in brown planthopper. **A.** Amino acid sequence of the maccessin precursor in brown planthopper. Yellow marker indicates signal peptide sequence. The six cysteines (blue marker) can form three disulfide bonds. The mature peptide is shown in gray background. The red-marked KR sites indicate cleavage sites for mature peptide. Note that there is no C-terminal amidation signal. The amino acids marked in red font indicate the peptide detected by mass spectrometry. **B.** The tissue distribution of *maccessin* was analyzed by semi-quantitative RT-PCR. Whole animals (female, male), male accessory glands (AG) and whole males with accessory glands removed (Male –AG), were assayed. **C.** The gene-silencing efficacy of *Maccessin* in male insects following dsRNA injection assayed by qPCR. The small circles denote the number of replicates. Data are shown as mean ± s.e.m. Student’s t-test. \*\*\**P* < 0.001. **D.** Receptivity of virgin and mated females, scored as the percentage of females that copulated within 30 min. The small circles denote the number of replicates; the numbers below the bars denote total number of animals. Data are shown as mean ± s.e.m. Mann–Whitney test. \**P* < 0.05, and ns (non-significant), *P* > 0.05. **E.** Number of eggs laid per female in 48 h. The dsRNA was injected in males at the end of the fifth instar, and the first mating took place after two days of eclosion. The numbers below the bars denote total number of animals. At least four biological replicates with at least five insects per replicate for each experiment. Data are shown as mean ± s.e.m. Student’s t-test. ns (non-significant), *P* > 0.05. **F.** Mating receptivity shown as percentage of virgins that copulated within 30 min and tested six hours after injection. Each virgin was injected with 30 nl of 1×PBS or 100 μmol/L maccessin. The small circles denote the number of replicates; the numbers below the bars denote total number of animals. Data are shown as mean ± s.e.m. Mann–Whitney test. \**P* < 0.05. **G.** Number of eggs laid after injection of maccessin in virgin females. Data are shown as mean ± s.e.m. Student’s *t*-test. \*\*\**P* < 0.001, \**P* < 0.05, and ns (not significant), *P* > 0.05, for comparisons against *dsgfp* injected in C-G. The numbers below the bars denote total number of animals. At least four biological replicates with at least five insects per replicate for each experiment. Each virgin was injected with 30 nl of 1×PBS or 100 μmol/L maccessin. **H.** Percentage of virgins laying eggs during 48 h after maccessin injection. The numbers below the bars denote total number of animals. At least four biological replicates with at least five insects per replicate for each experiment. Each virgin was injected with 30 nl of 1×PBS or 100 μmol/L maccessin. Chi-square test with the Yates’ correction ns (not significant), *P* > 0.05, for comparisons against PBS injection.

The *mac*c gene could not be detected in related insect species, such as small brown rice planthopper, *Laodelphax striatellus*, and whitebacked planthopper, *Sogatella furcifera* or other more distantly related insects including fruit flies and mosquitos (Figure 3-figure supplement 1A). However, we found a gene homologous to *mac*c in the closely related species *Nilaparvata muiri.* Thus, the macc peptide is well conserved between *Nilaparvata muiri* and *Nilaparvata lugens* (Figure 3-figure supplement 1B).

### The novel peptide maccessin reduces female receptivity but does not induce oviposition in *N. lugens*

Next, we asked whether the novel peptide macc plays a role in the PMR of BPHs. We diminished the expression of the *macc* gene in males by injection of dsRNA with a knockdown efficiency of more than 90% (Figure 3C), and found that a significant number of females that had been mated with these males would mate again with wild type males (Figure 3D). Such re-mating is never observed in the control (*dsgfp*-injected) group. However, the re-mating rate is small with only 7 percent of the females courted in the secondary mating being receptive (Figure 3D). The number of eggs laid per female displayed no difference between females mated with *dsgfp*- and *dsmacc*- injected males (Figure 3E).

If macc is transferred from a male to a female during copulation it could enforce his paternity by reducing receptivity of the female to further mating attempts. A similar response should be seen after macc peptide has been injected into a virgin female. To test this, we generated recombinant macc for injections. Next, we injected individual wild-type virgin females with either buffer or mature macc peptide and allowed them to recover for 6 hr in groups. This extended recovery time was required because virgins tested shortly after injection did not mate regardless of the substance injected. After recovery, injected females were exposed to wild-type males for 30 min, an exposure time that was sufficient for nearly all control females to show receptivity. We found that injection of macc significantly reduces virgin female receptivity (Figure 3F). Besides this, we observed an obvious post mating response in virgin females 6 hours after injection with macc, such as ovipositor extrusion, which is never seen in PBS injected females. However, injection of macc did not induce oviposition in virgin females (Figure 3G and H). In summary, our data show that females mated with males with diminished *macc* will mate again and that injection of macc in females reduces receptivity, but does not induce oviposition of virgin BPHs.

### An ITP-like peptide (ITPL-1) also reduces female receptivity but does not induce oviposition in *N. lugens*

Previous work identified another ITP precursor gene (Accession number: XP_039277955) in the BPH that gives rise to an amidated ITP peptide (ITPa) and four distinct ITPL transcripts (ITPL-1-4) containing different 5’ UTRs [48]. The open reading frame (ORF) and the 3’ UTR regions of the four ITPL transcripts are equivalent and encode identical non-amidated ITPL peptides (Figure 4-figure supplement 1) [35, 48].

**Figure 4.**
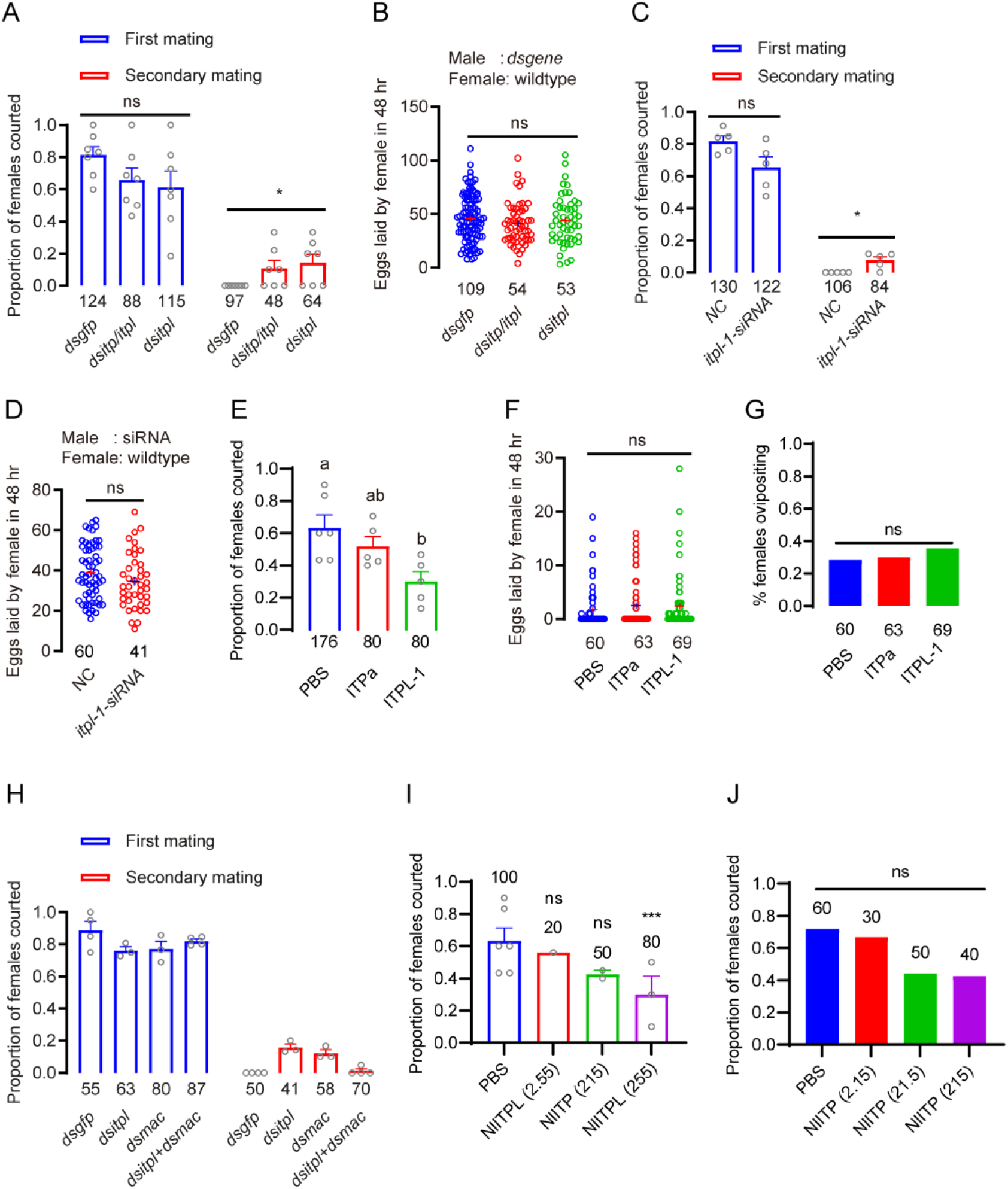
ITPL-1 also reduces female receptivity but not ovipostion in brown planthopper. **A.** Receptivity of virgin and mated females after *itp/itpl* and *itpl* knockdown by dsRNA injections, scored as the percentage of females that copulated within 30 min. The small circles denote the number of replicates; the numbers below the bars denote total number of animals. Data are shown as mean ± s.e.m. Mann–Whitney test, for comparisons against *dsgfp* injection control. \**P* < 0.05, and ns (non-significant), *P* > 0.05. **B.** Number of eggs laid per female in 48 h. dsRNA was injected in males at the end of the fifth instar, and the first mating took place two days after eclosion. The small circles and the numbers below the bars denote total number of animals. Data are shown as mean ± s.e.m. Student’s t-test. ns (not significant), *P* > 0.05, for comparisons against *dsgfp* injected. **C.** The mating receptivity rates of female to *NC (negative control)* and *Nlitpl1-siRNA* injected male courtship. The *Nlitpl1-siRNA* is designed to target a sequence specific to *Nlitpl1*, differentiating it from the other four spliceosome components. The small circles denote the number of replicates; the numbers below the bars denote total number of animals. Data are shown as mean ± s.e.m. Mann–Whitney test. \**P* < 0.05, and ns (non-significant), *P* > 0.05. **D.** Number of eggs laid per female in 48 h. The experimental protocol and symbols are the same as Fig. 4B. The small circles and the numbers below the bars denote total number of animals. Data are shown as mean ± s.e.m. Student’s *t*-test. ns (not significant), *P* > 0.05, for comparisons against *NC* injected. **E.** Mating receptivity shown as percentage of virgins that copulated within 30 min as tested six hours after injection. Each virgin was injected with 30 nl of 1×PBS , 400 μmol/L ITP or 400 μmol/L ITPL. The numbers below the bars denote total number of animals. Data are shown as mean ± s.e.m. Groups that share at least one letter are statistically indistinguishable; Kruskal–Wallis test followed by Dunn’s multiple comparisons test with *P* < 0.05. **F.** Number of eggs laid after injection of ITPa or ITPL-1 peptide in virgin females. The small circles and the numbers below the bars denote total number of animals. Data are shown as mean ± s.e.m. Student’s *t*-test. Ns (not significant), *P* > 0.05, for comparisons against PBS injected. **G.** Percentage of virgins laying eggs during 48 h after ITP or ITPL injection. The numbers below the bars denote total number of animals. Chi-square test with the Yates’ correction ns (not significant), *P* > 0.05, for comparisons against PBS injection.

The four ITPL transcripts display differential spatio-temporal expression patterns, where ITPL-1 is exclusively expressed in males, and specifically only in the male reproductive system [35]. However, the three other splice forms *itpl-2-*4 are expressed in other tissues in both males and females [35]. We confirmed these findings by RT-PCR and qPCR and found that *itpl-1* is exclusively expressed in the MAG and not in females (Figure 4-figure supplement 2A-C).

As noted above, the mature ITPL that can be generated from *itpl*-1 transcript is identical to the ones derived from itpl-2-4, suggesting that this peptide can be produced also in females. Nevertheless, we next asked whether male-derived ITPL-1 plays a role in the PMR of BPHs. Thus, we injected dsRNA that target *itp/itpl* (it is not possible to only silence *itp*) or only *itpl* (dsRNA to target exon 3) to determine whether peptide-deficient males can induce a PMR in female BPHs. Our data show that dsRNA significantly reduced the transcript levels of *itp* and the four splice forms *itpl1-4* in BPH (Figure 4-figure supplement 2D-G). Like for *macc*, we observed that a number of females who first had mated with *dsitp/itpl* and *dsitpl* males did remate with wildtype males, which does not occur after first mating with the *dsgfp* control males (Figure 4A). However, this remating rate is lower than 20 percent of the females courted in the secondary mating (Figure 4A). The number of eggs laid per female displayed no difference between females mated with *dsgfp*, *dsitp/itpl* and *dsitpl* injected males (Figure 4B). To specifically silence the *itpl-1* isoform, we synthesized small interfering RNAs (siRNAs), which target the exon 1a (Figure 4-figure supplement 1). We again observed that a number of females who first mated with itpl-1 siRNA males did remate with wild type males, which was not seen in controls (Figure 4C). However, the number of eggs laid was not changed after copulating with *itpl-1* siRNA males (Figure 4D). In summary, females who mated with *itpl-1* knockdown males displayed low rate of receptivity in the second mating (similar to the novel *macc*).

Next, we used recombinant amidated ITP (ITPa) and non-amidated ITPL-1 to test the effect of injection in females on the PMR. Individual wild-type virgin females were injected with either buffer, mature ITPa or ITPL-1 peptide. We found that injection of ITPL-1 reduced the receptivity of virgin females (Figure 4E). However, ITPa only has a weak effect on receptivity of virgin females (Figure 4E). We hypothesize that since ITPa is not produced in the MAG, the effect of injected peptide on female receptivity might reflect the action of endogenous female ITPa in post-mating physiology. Furthermore, since also peptides identical to ITPL-1 (ITPL-2-4) could be produced endogenously in females, we cannot exclude that also injected ITPL-1 mimics endogenous peptide, at least partly. We did not find that oviposition increased after ITPa and ITPL injection into virgin females (Figure 4F and G).

### Myoinhibitory peptides (MIPs) and *Drosophila* sex peptide (SP) do not trigger a post mating response in *N. lugens*

We found a seminal fluid-derived peptide, macc, that can trigger a PMR in the female BPH, but since it is an “orphan” peptide unrelated to previously known ones the receptor is unknown. Thus, we asked whether the MIPR previously implicated in *Drosophila* and other insects may act as a receptor of macc. However, first, we asked whether the known MIPR ligands SP or MIPs play any role in the PMR of BPHs. We thus tested whether *Drosophila* SP can induce a PMR in BPH. As a control, we showed that injection of SP significantly inhibits receptivity of virgin female *Drosophila* (Figure 5A). However, injection of *Drosophila* SP does not diminish receptivity of virgin BPHs (Figure 5B).

**Fig. 5.**
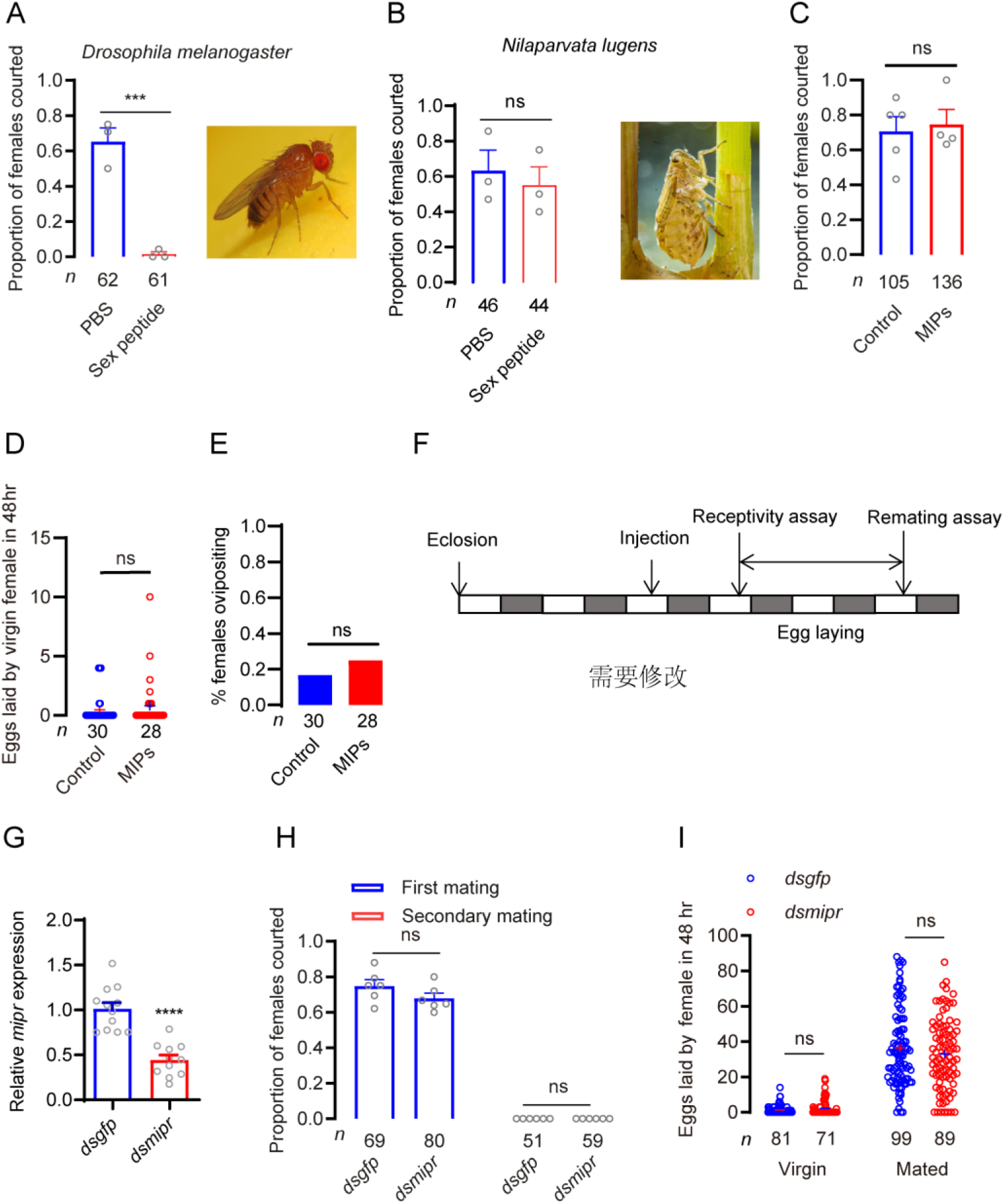
MIP and MIPR are not involved in the post-mating response of brown planthoppers. **A**. Mating receptivity shown as percentage of virgins that copulated within 30 min as tested six hours after sex peptide (SP) injection in *Drosophila melanogaster*. The small circles denote the number of replicates; the numbers below the bars denote total number of animals. Data are shown as mean ± s.e.m. Mann– Whitney test. \*\*\**P* < 0.001. **B**. Mating receptivity shown as percentage of virgins that copulated within 30 min as tested six hours after SP injection in *Nilaparvata lugens*. Each virgin was injected with 30 nl of 600 μmol/L SP. Data are shown as mean ± s.e.m. Mann–Whitney test. ns: no significant. **C**. MIP mixture injection does not affect female receptivity. Mating receptivity shown as percentage of virgins that copulated within 30 min as tested six hours after MIPs injection. Each virgin was injected with 30 nl of 1×PBS or 600 μmol/L MIPs. MIPs are a blend of MIP1, MIP2, MIP3 and MIP4. The small circles denote the number of replicates; the numbers below the bars denote total number of animals. Mann–Whitney test. ns: no significant. **D.** Number of eggs laid after injection of MIPs in virgin females. The small circles denote the total number of animals. Ns: no significant *P* > 0.05; Student’s t test. **E.** Percentage of virgins laying eggs during 48 h after MIP injection. The numbers below the bars denote total number of animals. Chi-square test with the Yates’ correction ns (not significant), *P* > 0.05. **F.** Protocol for behavioral experiments in G and H. **G.** Downregulation of *mipr* gene using *mipr*-RNAi leads to a reduction in mRNA expression level. \*\*\*\**P* < 0.0001; Student’s t test. The small circles denote the replicates. **H.** Receptivity of virgin and mated females, scored as the percentage of females that copulated within 30 min. Data are shown as mean ± s.e.m. Mann–Whitney test. Ns (not significant), *P* > 0.05. The small circles denote the number of replicates; the numbers below the bars denote total number of animals. **I.** Number of eggs laid per female in 48 h. The small circles denote the total number of animals. Data are shown as mean ± s.e.m. Student’s *t*-test. Ns (not significant), *P* > 0.05.

MIPs have been reported as the ancestral ligands of the promiscuous *Drosophila* SP receptor (also known as MIPR) [17, 18, 51, 52], but do not induce a PMR in *Drosophila* [17, 18]. Since MIP signaling is present ubiquitously in insects (<u>Kim et al.,</u> <u>2010</u>; <u>Poels et al., 2010</u>;) and the MIPR has been implicated in the PMR in a few insect species [see [19]], we asked whether MIP signaling is involved in the PMR in BPHs.

First, we cloned the *mip* gene of the BPH and found that it encodes four mature MIP peptides, MIP1 - MIP4 (Figure 5-figure supplement 1). Of these, MIP2 is predominantly expressed in BPH with eight paracopies in the precursor (Figure 5-figure supplement 1). Next, we examined the expression pattern of *mip* in BPHs. Investigating different developmental stages by real-time PCR, we found that *mip* transcript levels are boosted in third instar larvae and adult males. Transcripts are more abundant in adult males than in females. Of the different tissues, *mip* was detected in highest levels in the head of both male and female BPHs (Figure 5-figure supplement 2A and B).

We synthesized four mature MIP peptides for testing a possible role in the PMR in BPHs. For this test, we injected a mix of MIPs (MIP1 – MIP4) at different doses into the abdominal hemocoel of virgin females and allowed them to recover for 6 hr in groups. This extended recovery time was required because virgins tested shortly after injection did not mate regardless of the substance injected. After recovery, injected females were exposed to wild-type males for 30 min, an exposure time that was sufficient for nearly all control females to show receptivity. Similar to results in *Drosophila* [17, 18], we found that injecting a mix of MIPs into the abdominal hemocoel of virgin females does not decrease their receptivity (Figure 5C) or induce oviposition (Figure 5D and E).

### The MIP receptor is not involved in the post mating response of BPHs

The MIP receptor (MIPR) has been reported to be involved in the PMR of *Drosophila* [16], tobacco cutworm [25] and cotton bollworm [24]. However, a recent study indicated that MIPR is not required for refractoriness to remating or induction of egg laying in *Aedes aegypti* [53]. The MIPR has been identified in most of insect species, including BPHs [17, 18, 47] (Figure 5-figure supplement 3). The MIPR of BPHs is orthologous to the *Drosophila* MIPR (Figure 5-figure supplement 3). Hence, we asked whether the MIPR mediates the post-mating switch in BPH behavior. As a first step to address this question, we cloned the *mipr* gene of BPH (Figure 5-figure supplement 4). The MIPR of the BPH displays a typical seven transmembrane domain and it clusters with MIPR of other insect species (Figure 5-figure supplement 4A and B). We furthermore investigated the expression pattern of *mipr* in tissues of the BPH. The *mipr* is found throughout the nymphal and adult stages, and is expressed more predominantly in the head than other tissues (Figure 5-figure supplement 5A and B).

Next, we asked whether the MIPR plays a role in the PMR of BPHs using RNAi technique. The efficacy of the RNAi was tested by qPCR of whole animals and we found that the expression of *mipr* was significantly reduced (Figure 5G). We used a protocol in which individual virgin females (*dsgfp*- or controls that were *dsmipr*-injected) were first tested for receptivity towards a naive male (Figure 5F). Those females that mated were then allowed to lay eggs for 48 h before being retested for receptivity with a second naive male (Figure 5F). In the initial mating assays, virgin *mipr* RNAi females were as receptive as the control females (Figure 5H). When testing mated females for a second mating we did not detect any difference between controls and BPHs with *mipr* knockdown, suggesting that the MIPR has no effect on the refractoriness to remating (Figure 5H).

Next, we tested whether increased post-mating oviposition requires MIPR signaling by applying *mipr* RNAi. We found that both *dsgfp*-injected and *dsmipr*-injected females laid very few eggs if they were not mated (Figure 5I). We hypothesized that if the MIPR is required for post mating oviposition, *dsmipr*-injected females would lay few or no eggs even after mating. However, the number of eggs laid by mated *dsmipr*-injected females was not significantly different from that laid by those that were *dsgfp*-injected (Figure 5I). We thus conclude that neither MIPs nor the MIPR are required for increased post mating oviposition in *N. lugens*, and the MIPR is not likely to be the receptor of the novel peptide macc.

## Discussion

A mating-induced switch in female behavior and physiology to ensure a numerous and viable offspring, as well as to secure paternity, is common in insects, but only in *Drosophila melanogaster* the underlying mechanisms have been clarified in detail and the MAG-derived peptide SP identified as the main secreted signal [3, 4, 6, 7, 9, 11–13].

Another closely related MAG-derived peptide, DUP99B, is also contributing to the PMR in *D. melanogaster* [54, 55]. Since SP and DUP99B can be found only in a few *Drosophila* species, we set out to identify factors in the MAG of the brown planthopper (BPH) that might play a role similar to SP in inducing a PMR. First we established that BPH females display a distinct change in behavior and physiology after mating. This is manifested in a reduction in receptivity to mating males and an increase in ovulation lasting four days. Next, we showed that extract from MAG injected into female BPHs induced a significant decrease in receptivity, lasting about 24 h, and a significant increase oviposition. Then we asked what the active factor in MAG extract that induces the PMR might be. We therefore went on to perform a transcriptional analysis of MAG extract. As expected SP and MIPs were not detected, but one splice form of ion transport peptide, ITPL, and a novel 51 amino acid peptide were identified. A fragment of the latter was also detected by mass spectrometry. While the ITPL was identified also in other tissues of both males and females, (see also [35, 48]) we focused on the novel peptide, maccessin, which is male specific and only found in the MAG.

Virgin female BPHs mated to males with *macc* knockdown do not display a repression of the propensity to re-mate, whereas injections of recombinant macc peptide into virgin females render them less receptive to courting males. However, oviposition was not affected by these manipulations. Thus, we propose that *macc* mediates a male signal transferred in seminal fluid to reduce female receptivity to further courting males. In our experiments this signal only mediates the receptivity part of the total PMR seen after regular mating and the duration of the effect is shorter (only 24 h instead of 4 d). We speculate that the partial and shorter PMR effect seen in our macc experiments could be for the following reasons: (1) in *Drosophila* SP to exert its full effect over about a week needs to be bound to sperm when transferred to the female and thereafter gradually released [46], and thus macc injection without sperm may be less efficient and the peptide exposed to protease degradation, (2) it is likely that macc is not the only MAG-secreted factor required for a full PMR effect and therefore *macc* RNAi in males is not sufficient. In fact, we did identify another peptide, ITPL-1 in the MAG of BPHs and found that it also induces a partial effect on the female PMR.

Since the receptor of SP in *Drosophila* [16] can be activated also by MIPs [8, 17, 18], and the MIPR was implicated in the PMR in some insects [25] [24], we tested whether SP, MIPs or their receptor (MIPR) affects the PMR in BPHs. We found no effect of manipulating MIP signaling or injections of *Drosophila* SP and planthopper MIPs. The outcome of the MIPR knockdown experiment also suggests that this receptor is not required for macc signaling. Thus, since macc is a novel peptide with no sequence relation to previously identified peptides, its receptor is still unknown, and we were unable to manipulate this part of the signal pathway in female tissues for further tests.

It is intriguing that SP and Dup99B are found only in the genomes of a few *Drosophila* species related to *D. melanogaster* [15, 17, 56]. Furthermore, other peptides regulating a PMR response have not yet been unequivocally identified in any other insect, except the mosquito *Aedes aegypti* where post-mating receptivity to males was found affected by a MAG-derived peptide, HP-1 [27], as detailed below.

Since SP acts on the MIPR several studies investigated the involvement of MIP signaling in a PMR in various insects. However, MIP activated MIPR signaling seems not to regulate PMR in the insects studied (including the present study), although the MIPR and some unidentified ligand affects oviposition and ovary development in some insects [20–22, 25, 53]. In mosquitos a post-mating decline in female receptivity to further mating attempts is mediated by the MAG-derived head peptide, HP-1 (a form of short neuropeptide F, sNPF) and the sNPF receptor NPYLR1 [27]. However, the HP-1 signaling does not affect fecundity, host-seeking or blood-feeding. This is similar to the BPH where *macc* is MAG-derived signal that reduces female receptivity, but not fecundity. Thus, mosquito HP-1 and BPH macc may be partial functional analogs of SP.

What SP, HP-1 and macc have in common is that they are MAG-derived peptides that appear to be restricted phylogenetically. We found a macc precursor transcript only in the genomes of *Nilaparvata lugens* and *Nilaparvata muiri,* but not in the related small brown rice planthopper, *Laodelphax striatellus*, or the white-backed planthopper, *Sogatella furcifera,* or other more distantly related insects. Apparently MAG-derived secretory peptides undergo rapid evolution in certain species and in *Drosophila* SP repurposes an already existing receptor (MIPR) for distantly related MIP neuropeptides [15, 17, 18, 56]. Similarly, *Aedes* HP-1, has adopted an sNPF receptor [27]. Thus, SP displays some sequence similarities to MIPs [17, 18] and HP-1 is sNPF-like, while the macc sequence is unique making it a more complex task to select known GPCRs (or orphan receptors) for screening. The role of ITPL-1 needs to be further investigated. Since it also seems to be produced in female BPHs from the other splice variants itpl-2-4 it may act both via transfer from males at copulation and as an endogenously secreted peptides in females. Receptors for ITPa and ITPL peptides have been identified in the moth *Bombyx mori* and the fly *Drosophila* [57, 58]. Thus, receptor knockdown could be attempted in females for tests of PMR in future.

In summary, we identified a novel peptide, macc, in the MAG of BPHs that induces a PMR in mated females rendering them less receptive to further mating attempts. It remains to identify a receptor for this novel peptide and to characterize target circuits in the central nervous system that modulate the female behavior. Additionally, the role of ITPL-1 in the PMR should be further investigated, including the possible role of endogenous female ITPL-1 in regulating reproductive behavior and physiology, and a search for additional factors that ensures the fecundity in mated females and leads to a more complete PMR resembling that seen after mating.

## Materials and methods

### Experimental insects and husbandry

The brown planthopper *N. lugens* was reared on ‘Taichung Native 1’ (TN1) rice (*Oryza sativa* L.) seedlings in the laboratory and maintained at 27 ± 1 ^∘^C, with 70 ± 10% relative humidity, under a 16 h: 8 h light dark photoperiod [59]. The brown planthopper sensitive strain was originally supplied in 1995 by Zhejiang Chemical Technology Group Co., LTD.

### Extraction of male accessory glands proteins

300 pairs of male accessory glands were dissected in phosphate buffer at pH 7.2 and were transferred into 100 μl of 80% methyl alcohol on the ice. These samples were treated by ultrasonic homogenization. After centrifugation supernatants were collected and the precipitate washed twice with 80% methanol. The supernatant was mixed and freeze dried in a Speed Vac Vacuum concentrator.

### Mass spectrometry

#### Total protein extraction

We dissected the male accessory glands of brown planthoppers in PBS buffer, and collected 300 glands for each replicate (for a total of three biological replicates). The male accessory glands were quickly frozen in liquid nitrogen, ground into powder at low temperature, and quickly transferred it to a centrifuge tube pre-cooled with liquid nitrogen. We added an appropriate amount of protein lysis buffer (100 mM ammonium bicarbonate, 8M urea, 0.2% SDS, pH=8), mixed well and sonicated in an ice-water bath for 5 minutes to ensure complete lysis. Centrifugation was performed at 4°C and 12000 g for 15 minutes, and the supernatant collected. A final concentration of 10 mM DTT was added to the supernatant and reacted at 56°C for 1 hour. After that, an adequate amount of IAM was added and reacted at room temperature in the dark for 1 hour. Four times the volume of pre-cooled acetone was added at -20°C to precipitate for at least 2 hours at -20°C, then centrifugated at 4°C and 12000 g for 15 minutes to collect the precipitate. The precipitate was resuspended and washed with 1 mL of pre-cooled acetone at -20°C, centrifuge<u>d</u> at 4°C and 12000 g for 15 minutes, and the precipitate collected. The precipitate was air-dried, and then the protein precipitate dissolved by adding an appropriate amount of protein solubilization buffer (6M urea, 100 mM TEAB, pH=8.5).

#### Protein quality inspection

We used the Bradford protein assay kit to prepare a BSA standard protein solution according to the instructions, with a concentration gradient ranging from 0 to 0.5 µg/µL. We took different concentrations of the BSA standard protein solution and different dilutions of the test sample solution and added them to a 96-well plate, making up the volume to 20 µL, with each gradient repeated 3 times. We quickly added 180 µL of G250 staining solution, let it stand at room temperature for 5 minutes, and measured the absorbance at 595 nm. We plotted a standard curve using the absorbance of the standard protein solution and calculated the protein concentration of the test samples. We took 20 µg of protein test samples for 12% SDS-PAGE gel electrophoresis, with the conditions for the stacking gel electrophoresis being 80 V for 20 minutes, and the conditions for the separating gel electrophoresis being 120 V for 90 minutes. After the electrophoresis is completed, we stained with Coomassie Brilliant Blue R-250, and destained until the bands are clear.

#### Proteolysis

We took 120 µg of protein samples, added protein solubilization buffer to make up the volume to 100 µL, added 1.5 µg of trypsin and 500 µL of 100 mM TEAB buffer, mixed well, and digested at 37°C for 4 hours. We then added 1.5 µg of trypsin and CaCl2 for overnight digestion. We adjusted the pH to less than 3 with formic acid, mixed well, and centrifuged at room temperature at 12000 g for 5 minutes, took the supernatant, and slowly passed it through a C18 desalting column. After that, we used a washing solution (0.1% formic acid, 3% acetonitrile) to wash continuously three times, then added an appropriate amount of elution solution (0.1% formic acid, 70% acetonitrile), collected the filtrate, and freeze-dried it.

#### Liquid quality detection

We prepared mobile phase A solution (100% water, 0.1% formic acid) and B solution (80% acetonitrile, 0.1% formic acid). We dissolved the freeze-dried powder with 10 µL of A solution, centrifuged at 4°C at 14000 g for 20 minutes, and took the supernatant for sample injection, with 1 µg of sample for liquid chromatography-mass spectrometry (LC-MS) analysis. We used the EASY-nLC^TM^ 1200 nanoflow UHPLC system, equipped with a homemade pre-column (2 cm × 75 µm, 3 µm) and a homemade analytical column (15 cm × 150 µm, 1.9 µm). The liquid chromatography elution conditions were as depicted in Table S4. We employed the Q Exactive^TM^ HF-X mass spectrometer with a Nanospray Flex™ (ESI) ion source, set the ion spray voltage to 2.3 kV, and the ion transfer tube temperature to 320°C. The mass spectrometer operated in data-dependent acquisition mode, with a full scan range of m/z 350-1500, a primary mass spectrometry resolution set to 60000 (at 200 m/z), a maximum C-trap capacity of 3×10^6^, and a maximum injection time of 20 ms. We selected the top 40 parent ions with the highest intensity in the full scan for fragmentation using high-energy collision dissociation (HCD) for secondary mass spectrometry analysis. We set the secondary mass spectrometry resolution to 15000 (at 200 m/z), a maximum C-trap capacity of 1×10^5^, and a maximum injection time of 45 ms. The peptide fragmentation collision energy was set to 27%, the threshold intensity was set to 2.2×10^4^, and the dynamic exclusion window was set to 20 seconds. We generated raw mass spectrometry data (.raw).

#### Data analysis

We used the brown planthopper protein database to search all the result spectra with the search software Proteome Discoverer 2.2 (PD2.2, Thermo). We set the search parameters as follows: the mass tolerance for precursor ions was 10 ppm, and the mass tolerance for fragment ions was 0.02 Da. The fixed modification was carbamidomethylation of cysteine, the variable modification was oxidation of methionine, and the N-terminus was acetylated. We allowed up to 2 missed cleavage sites.

We enhanced the quality of the analytical results by further filtering the search results using the PD2.2 software: Peptide Spectrum Matches (PSMs) with a confidence level above 99% were considered reliable PSMs, and proteins that contained at least one unique peptide were considered reliable proteins. We retained only the reliable PSMs and proteins, and performed a False Discovery Rate (FDR) validation to remove peptides and proteins with an FDR greater than 1%.

We used the InterProScan software for GO and IPR functional annotation, which included databases such as Pfam, PRINTS, ProDom, SMART, ProSite, and PANTHER. We performed functional protein family and pathway analysis on the identified proteins using COG and KEGG. We conducted volcano plot analysis, clustering heatmap analysis, and pathway enrichment analysis for GO, IPR, and KEGG on Differentially Expressed Proteins (DPE). Additionally, we predicted potential protein-protein interactions using the STRING DB software (http://STRING.embl.de/).

### Gene cloning and sequence analysis

We used the NCBI database and BLAST programs for sequence alignment and analysis. Then we used EditSeq to predict Open Reading Frames (orfs). The primers were designed by tools in NCBI. According to the manufacturer’s instructions, total RNA was extracted by TRIzol reagents (Invitrogen, Carlsbad, CA, USA). We used HiScript III RT SuperMix for qPCR (+gDNA wiper) (Vazyme, Nanjing, China) reverse transcription kit to synthesize cDNA templates for cloning, and stored the synthesized cDNA templates at -20℃.

We predicted protein transmembrane fragments and topological structures through TMHMM v2.0 (http://www.cbs.dtu.dk/services/TMHMM-2.0/) (Krough et al., 2001). Multiple alignments on the complete amino acid sequences were performed using ClustalX (http://www.clustal.org/clustal2/). The phylogenetic tree was constructed using MEGA 10.0 software and the Maximum Likelihood Method, with 1000 repeated starts.

### Gene expression profile analysis

For the stage-specific expression study of *mip* and *mipr*, total RNA were extracted from pools of thirty individuals from the following developmental stages: 1^st^ to 5^th^ instar nymphs, adult male and female insects. For the tissue-specific expression study of *mip* and *mipr*, total RNA was isolated from various tissues including head, thorax and abdomen of three-day-old male adults, and head, thorax and abdomen of three-day-old virgin female adults. For conducting a tissue-specific expression analysis of *maccessin*, total RNA was extracted from pooled samples of multiple individuals across the following phases: virgin females and males three days post-emergence, male accessory glands, and male bodies with removed accessory glands. All samples were extracted by using TRIzol reagent (Invitrogen).

### Quantitative RT-PCR

The first-strand cDNA was synthesized with HiScript® II Q RT SuperMix for qPCR (+gDNA wiper) kit (Vazyme, Nanjing, China) using an oligo(dT)18 primer and 500 ng total RNA template in a 10 μl reaction, following the instructions. Real-time qPCRs of the various samples used the UltraSYBR Mixture (with ROX) Kit (CWBIO, Beijing, China). The PCR was performed in 20 μl mixture including 4 μl of 10-fold diluted cDNA, 1μl of each primer (10 μM), 10 μl 2 × UltraSYBR Mixture, and 6 μl RNase-free water. The PCR conditions used were as follows: initial incubation at 95°C for 10 min, followed by 40 cycles of 95°C for 10 s and 60°C for 45 s. *N. lugens* 18S rRNA was used as an internal control. Relative quantification was performed via the comparative 2^−△△CT^ method [60].

### RNA interference

For lab-synthesized dsRNA, *gfp, mipr, itp, itp/itpl* and *Maccessin* fragments were amplified by PCR using specific primers conjugated with the T7 RNA polymerase promoter (primers listed in Supplementary Table S3). The dsRNA was synthesized by a kit (MEGAscript T7 transcription kit, Ambion) according to the manufacturer’s instructions. The integrity and quantity of the double-stranded RNA (dsRNA) products were confirmed using 1% agarose gel electrophoresis and a Nanodrop 1000 spectrophotometer. Subsequently, the samples were stored at -70°C until further use. In order to achieve the effect of silencing target genes, 5 μg/μl dsRNA was injected into brown planthopper, male 40nl, female 50nl, and control group the same amount of *dsgfp*. Total RNA was individually collected from each insect on the day after reception assay, followed by extraction. The efficiency of gene silencing was subsequently assessed through qPCR.

### Peptide synthesis

Peptides were synthesized by Genscript (Nanjing, China) Co., Ltd. Myoinhibitory Peptides and Sex Peptide mass was confirmed by MS and the amount of peptide was quantified by amino acid analysis. Ion transport peptides and maccessin peptide were immune recombinant proteins expressed in CHO cell. These proteins were purified by AmMagTM Ni Magnetic Beads. The amino acid sequence of the peptides used in this study are: *N. lugens* Myoinhibitory Peptide 1: (MIP1): AWRDLQSSWamide; Myoinhibitory Peptide 2: (MIP2): GWQDMPSSGWamide; Myoinhibitory Peptide 3 (MIP3): GWQDLQGGWamide; Myoinhibitory Peptide 4 (MIP2): AWSSLRGTWamide; *D. melanogaster* Sex Peptide: (SP): WEWPWNRK{Hyp}TKF{Hyp}I{Hyp}S{Hyp}N{Hyp}RDKWCRLNLGPAWGGRC.

Mature proteins ITPa (comprising amino acids 23-113), ITPL (comprising amino acids 23-117), and maccessin (comprising amino acids 20-91), each fused with a 6xHis tag at their C-termini (designated as protein-His tag), were expressed in Chinese Hamster Ovary (CHO) cells. These proteins were subsequently purified using AmMagTM Ni Magnetic Beads. Additionally, the mature ITPa protein with an amide modification at the C-terminus (referred to as ITPa-amide) was also expressed in CHO cells. All these proteins were codon-optimized for expression in mammalian cells.

### Behaviour assays

#### 1. Post mating response

For first mating assay, a couple of virgin females and males (3 days after eclosion) were kept for 30 min in a 24 mm (diameter) × 95 (height) mm transparent circular tube with rice seedlings. The mated females were subjected to re-mating assay every 24 hours for 1-5 days after first mating, while the virgin females of the same age were used as controls.

For egg laying, virgin or mated females (3 days after eclosion) were kept in a 24 mm × 95 mm transparent circular tube with rice seedling. Each female was numbered and moved into a new tube every 24 hours. The number of eggs in the rice seedlings was counted.

#### 2. Tests if SFPs induce a post-mating response

For experiments to test SFPs-injected females, virgin females that had eclosed 3 days earlier were selected for injection. Reception assays were performed with males of the same age placed in a single pair in the tube 3h, 6h, 12h, 24h and 36h after injection. Each female was injected with the equivalent of half of accessory gland.

On the day after eclosion, each virgin was injected with 30 nl SFPs or solvent and placed in the transparent circular tube with rice seedling to lay eggs for 24 h.

#### 3. Peptide injection to test virgin receptivity

For experiments using peptide-injected females (3 days after eclosion), the mating experiment was conducted 6 hours after injection, when females had fully recovered from the wound.

On the day after eclosion, each virgin female was administered an intra-abdominal injection of 30 nl of mature peptide or PBS. Subsequently, they were housed in a transparent cylindrical tube with a rice seedling, where they were allowed to oviposit for a period of 48 hours.

#### 4. Effect of silencing female *mipr* gene on post-mating response

For the effect of silencing the female brown planthopper *mipr* gene on the post-mating response, the experimental protocol is shown in Figure 2D. After recovering for 1 day, the virgin females were mated with wild type males. The mated females were re-mated with virgin males 2 days after the first reception assay. Between the first reception and the re-mating assay, mated females were placed in tubes with rice seedlings to lay eggs.

#### 5. Effect of silencing the male *maccessin* gene on post-mating response

As for experiments using dsRNA-injected males, co-caging with female virgins of the same age was 3-4 days after injection, when the *maccessin* gene was silenced to the greatest extent. Mated females were collected and remated with wild-type virgin males 2 days later. Between the reception and remating assays, the mated females were introduced into tubes with rice seedlings to facilitate oviposition.

Following the daily activity of the brown planthopper, the mating experiments were scheduled between 3 p.m. to 7 p.m. The male and female insects were kept in the tubes for half an hour to see if mating took place. Each experiment was repeated no less than 3 times, with at least 10 insects per repetition.

### RNA-seq analysis

Total RNA from 150 virgin male accessory glands was isolated three days post-emergence using TRIzol reagent (Invitrogen), following the manufacturer’s protocol. Library construction and sequencing was performed by Novogene with Illumina HiSeq2000 platform (Novogene Bioinformatics Technology Co.Ltd, Beijing, China).

After filtering out low-quality sequences, the raw data were subjected to analysis. Sequence alignment was conducted against the *Nilaparvata lugens* genome, accessible via the NCBI database (https://www.ncbi.nlm.nih.gov/genome/?term=Nilaparvata+lugens), using Hisat2 v2.0.5. The gene expression levels derived from RNA sequencing data were normalized using the FPKM method, which accounts for both sequencing depth and gene length in the calculation of read counts, making it a widely adopted approach for estimating gene expression levels. Differential gene expression analysis was executed with the DESeq2 R package (version 1.16.1). Subsequently, Gene Ontology (GO) enrichment and KEGG pathway analyses were performed using the clusterProfiler R package.

### Proteome analysis

The tissues were isolated from 150 virgin male accessory glands three days post-emergence. The label-free quantitative method involves mass spectrometry analysis of enzymatically digested proteins. Utilizing raw mass spectrometry data, a search was conducted against the RNA-seq database to identify proteins based on the search outcomes. Subsequently, an association analysis with the transcriptome data was conducted to elucidate the relationships between protein expression and gene sequences.

### Statistics

We employed GraphPad Prism 9 software for data visualization and statistical analysis. Data presented in this study were first verified for normal distribution by D’Agostino– Pearson normality test. If normally distributed, Student’s *t* test was used for pairwise comparisons, and one-way ANOVA was used for comparisons among multiple groups, followed by Tukey’s multiple comparisons. If not normally distributed, Mann–Whitney test was used for pairwise comparisons, and Kruskal–Wallis test was used for comparisons among multiple groups, followed by Dunn’s multiple comparisons. All data are presented as mean ± s.e.m. All data are collected from at least four independent experiments. Every independent experiment used at least five insects.

## Supporting information

All of supplemetal data

## ACKNOWLEDGMENTS

This research was supported by the National Key Research and Development Program of China (2022YFD1700200), the National Natural Science Foundation of China (No. 32472542), the Guidance Foundation of the Sanya Institute of Nanjing Agricultural University (NAUSYMS15) and the Fundamental Research Funds for the Central Universities (No. KJJQ2024016). We thank Dr. Mariana Wolfner for commenting on an earlier version of this paper.

## Supporting information Author Contributions

Conceptualization: Shun-Fan Wu, Dick R. Nässel. Data curation: Yi-Jie Zhang, Ning Zhang, Ruo-Tong Bu, Shun-Fan Wu. Formal analysis: Yi-Jie Zhang, Shun-Fan Wu. Funding acquisition: Shun-Fan Wu, Cong-Fen Gao. Investigation: Yi-Jie Zhang, Ning Zhang, Ruo-Tong Bu. Supervision: Shun-Fan Wu, Cong-Fen Gao, Dick R. Nässel. Validation: Yi-Jie Zhang, Ning Zhang, Ruo-Tong Bu. Writing – original draft: Yi-Jie Zhang, Shun-Fan Wu, Dick R. Nässel. Writing – review & editing: Shun-Fan Wu, Dick R. Nässel.

---

**Figure 1 - figure supplement 1.**
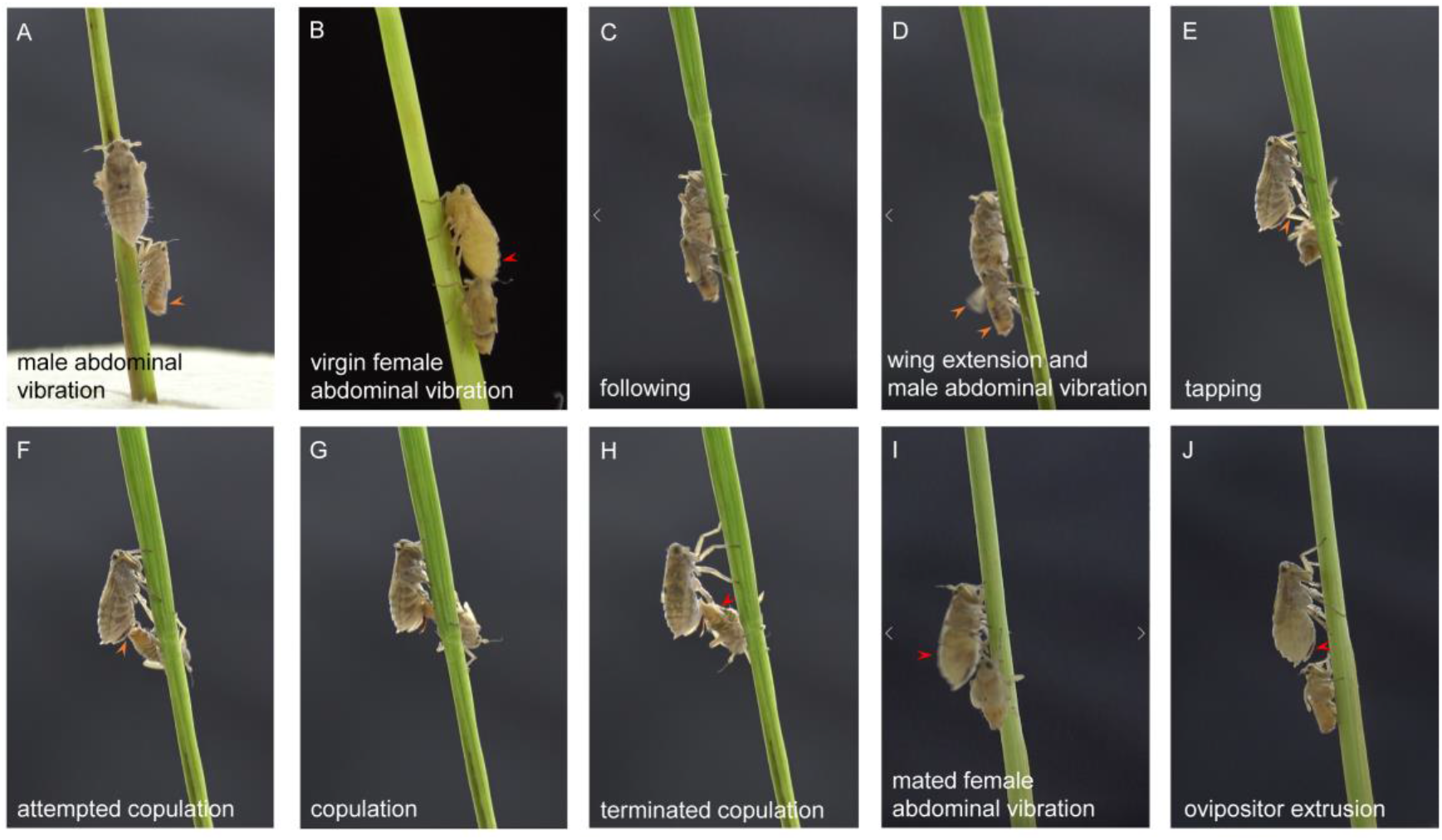
**Mating and post-mating behavior of brown planthoppers. A-H**: The mating behavior sequence includes eight steps (A-H, following, wing extension, abdominal vibration, abdominal rubbing, attempted copulation, copulation, terminated copulation and leaving). **I** and **J**: The post-mating response includes two behaviors, female abdominal vibration and ovipositor extrusion. The bigger insect is the female and smaller is the male. **Video S1** Video S1. Courtship behavior of brown planthopper. Video S2 **Video S2**. Post-mating behavior of brown planthopper. Video S3 **Video S3. Post-mating behavior of virgin brown planthopper after injection with seminal fluid proteins (SFPs).**

**Figure 2-figure supplement 1.**
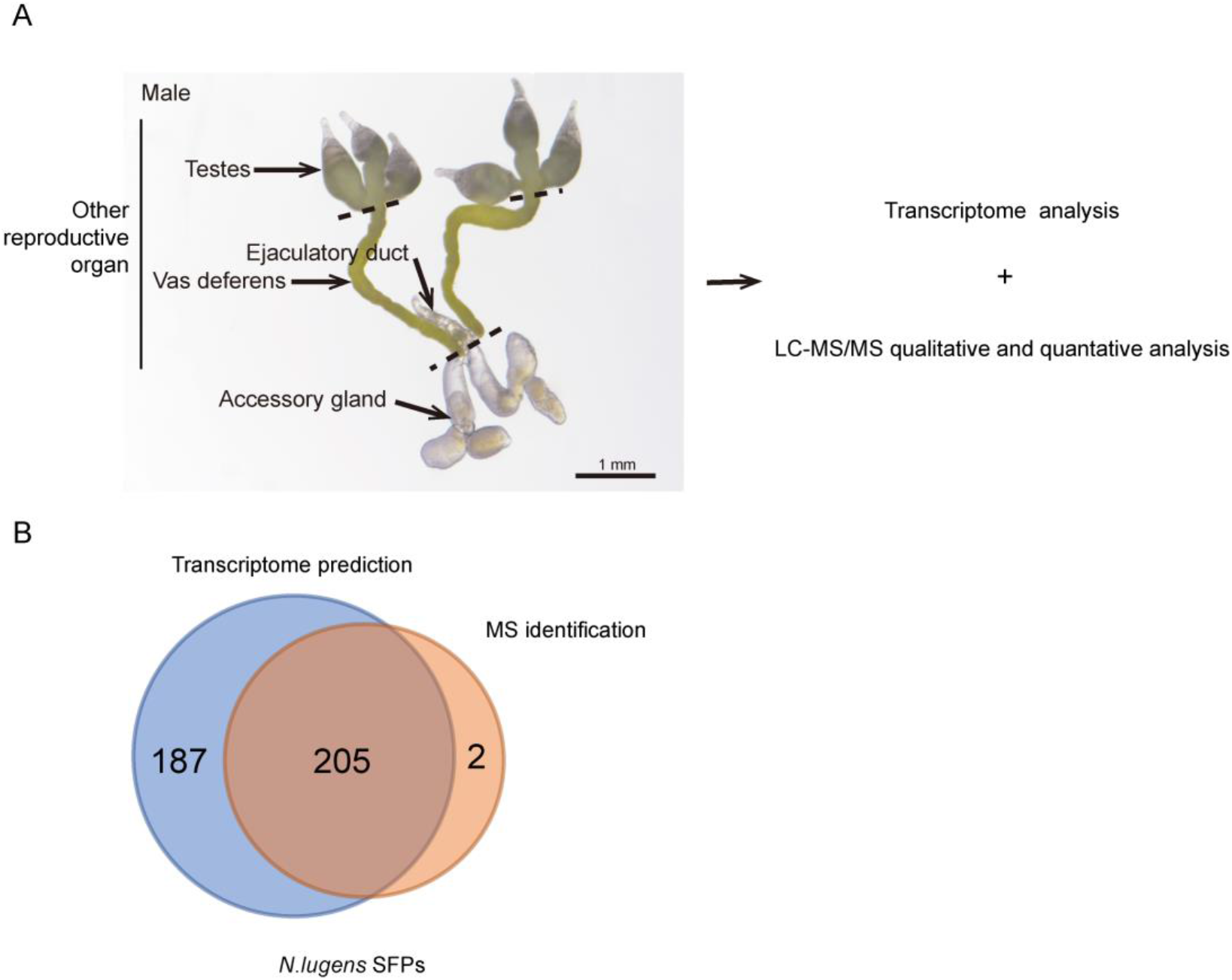
(A) Workflow for identification and quantitation of seminal fluid proteins in the brown planthopper, *N. lugens*. Dissected accessory glands were used for extraction and subjected to transcriptome and proteome analysis using liquid chromatography (LC) and mass spectrometry (MS/MS). (B) Venn diagram of the numbers of predicted seminal fluid proteins comparing transcriptome prediction and MS identification.

**Figure 2-figure supplement 2.**
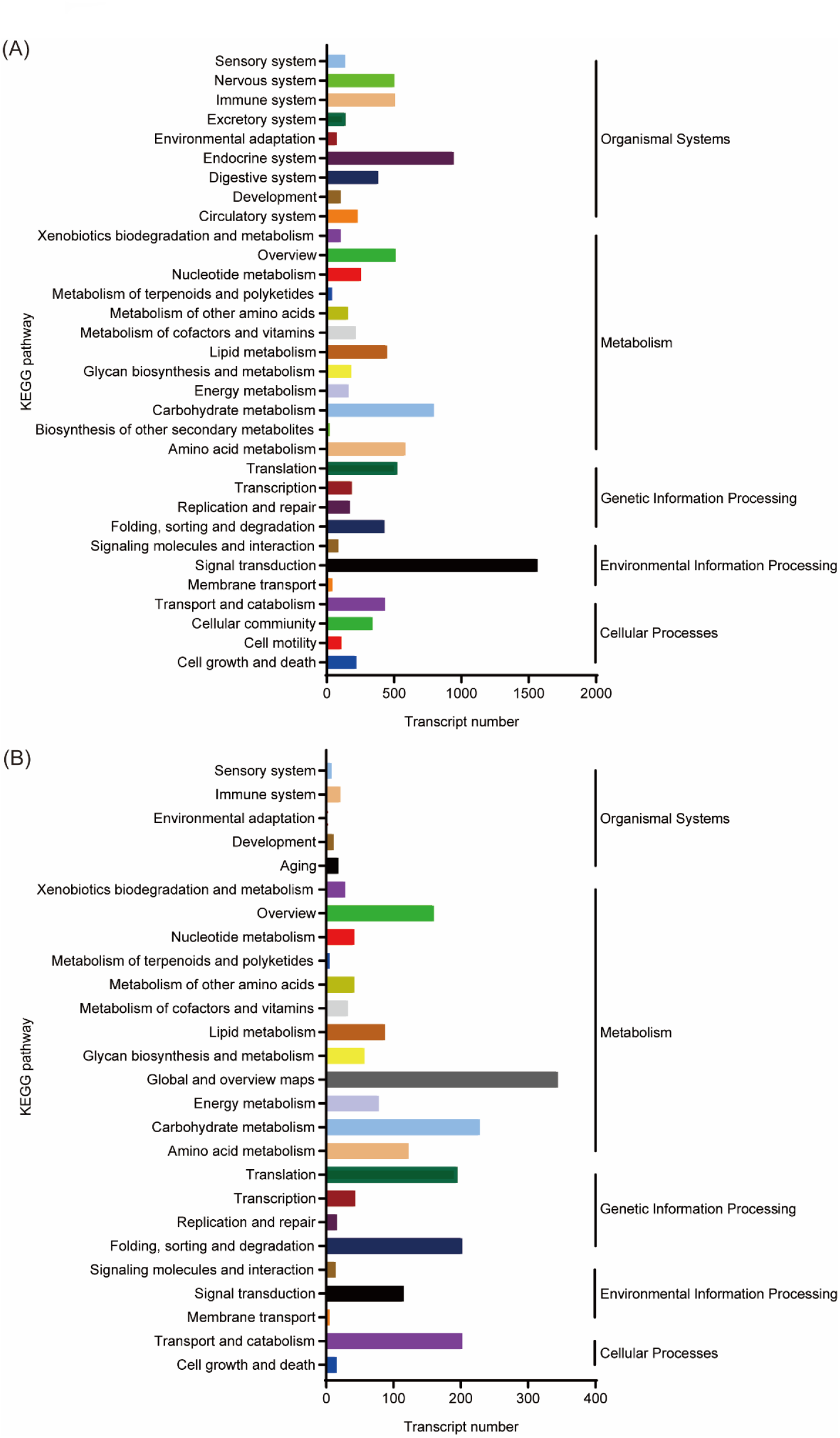
Pathway assignment based on KEGG analysis. (A) Classification based on transcript. (B) Classification based on protein.

**Figure 3 - figure supplement 1.**
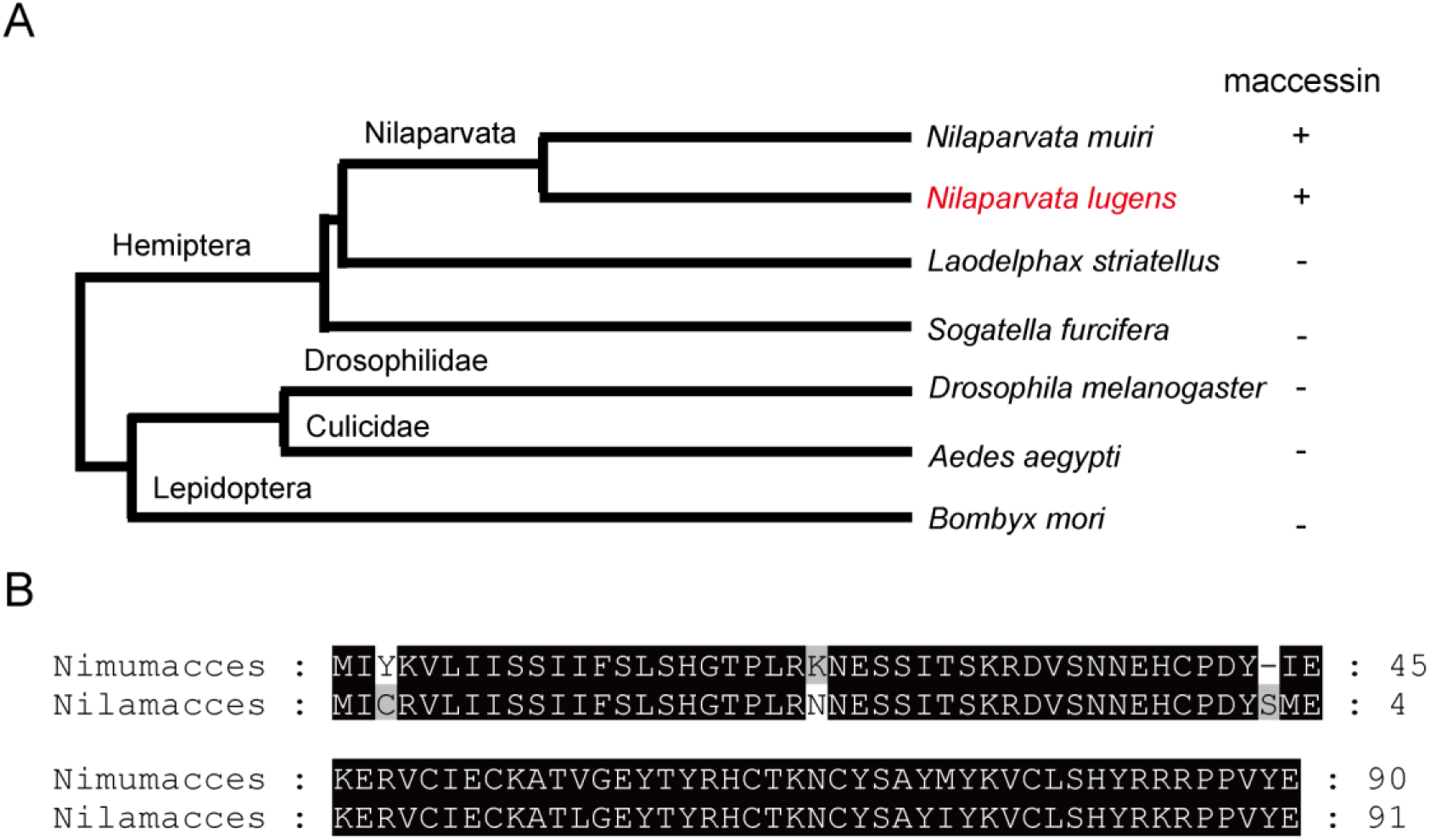
(A) A gene encoding maccessin precursors was detected in *Nilaparvata muiri* and *Nilaparvata lugens,* but not present in other planthoppers, or other insects such as fruit flies, mosquito and silkworm. (B) The alignment of maccessin protein sequence between *Nilaparvata muiri* (upper) and *Nilaparvata lugens* (lower).

**Figure 4 - figure supplement 1.**
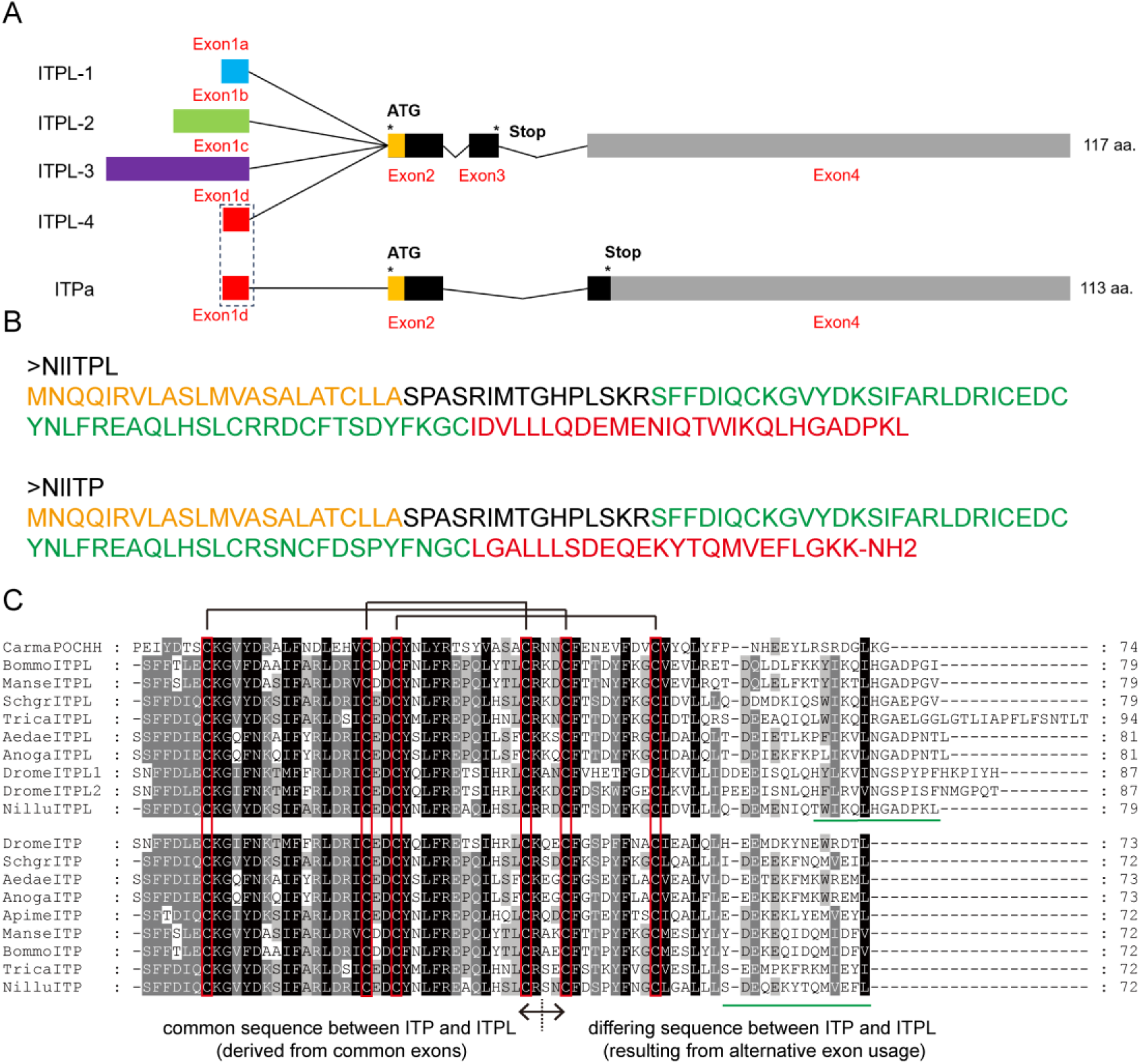
Sequence analysis of ITP/ITPL. **A.** The identified ion transport peptide/ ITP-like (ITPa/ITPL) transcripts. The coloured blocks represent exons within the *N. lugens* ITP/ITPL transcripts. Exons 1a, 1b, 1c and 1d are alternative 5’ untranslated regions used by ITP and ITPLs. * denote the start codon and stop codons of the transcripts. **B.** Amino acid sequence of ITP and ITPL of brown planthopper. Orange indicates sequence of signal peptide; green indicates mature peptide sequence; red indicates difference sequence of ITP and ITPL. Note that four slice forms of *itpl* are known (*itpl-1-4*), which all could give rise to the same mature ITPL peptide. **C.** Multiple comparisons of ITP and ITPL mature peptides in brown planthopper and other species. The red frames indicate conserved cysteines. Deduced ITP and ITPL sequences are shown for *Manduca sexta* (Manse, AY950500, AY950501), *Bombyx mori* (Bommo, AY950502, AY950503), *Schistocerca gregaria (Schgr)*, *Apis mellifera (Apime)*, *Aedes aegypti* (AY950504, AY950505, AY950506), *Anopheles gambiae (Anoga)*, *Drosophila melanogaster (Drome)*, and *Tribolium castaneum* (Trica, EFA07585).

**Figure 4-figure supplement 2.**
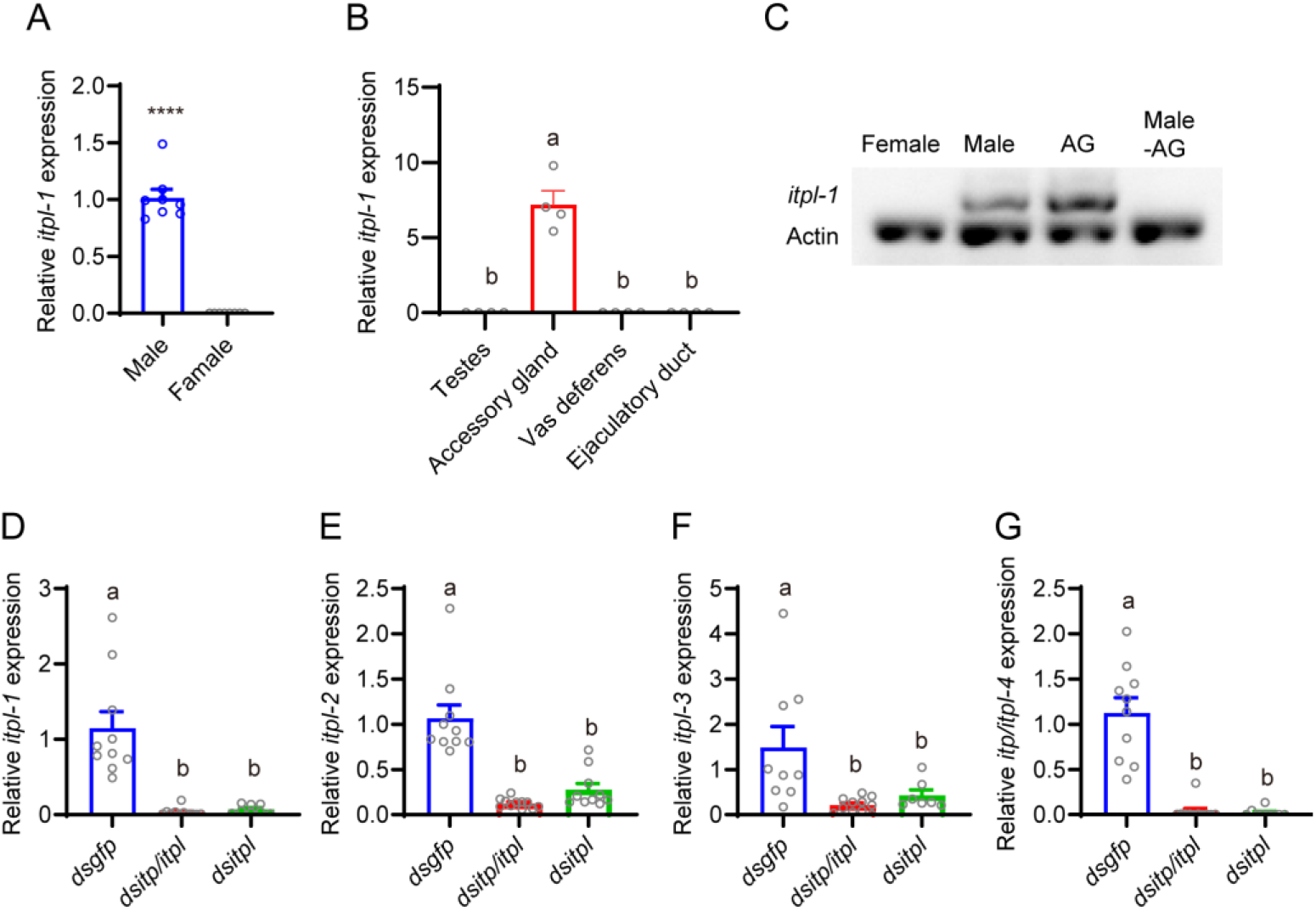
Distribution of *itpl-1* transcript in different genders and tissues of brown planthopper. **A.** Relative expression of *itpl-1* gene in *N. lugens* at different genders. Data are shown as mean ± s.e.m. Student’s *t*-test. ****, *P* < 0.0001, for comparisons against *dsgfp* injected. **B.** Relative expression of *itpl-1* gene in *N. lugens* at different tissues in male reproductive system. Data are shown as mean ± s.e.m. Groups that share at least one letter are statistically indistinguishable; Kruskal–Wallis test followed by Dunn’s multiple comparisons test with *P* < 0.05. **C.** The tissue distribution of *itpl-1* was analyzed by semi-quantitative RT-PCR. RNA samples from adult females, adult males, male accessary gland (AG) alone and adult male without accessary gland (male -AG). **D-G.** Relative expression of different spliceosomes of *itpl-1-4* gene in males injected with dsRNA. Data are shown as mean ± s.e.m. Groups that share at least one letter are statistically indistinguishable; Kruskal–Wallis test followed by Dunn’s multiple comparisons test with *P* < 0.05.

**Figure 5 - figure supplement 1.**
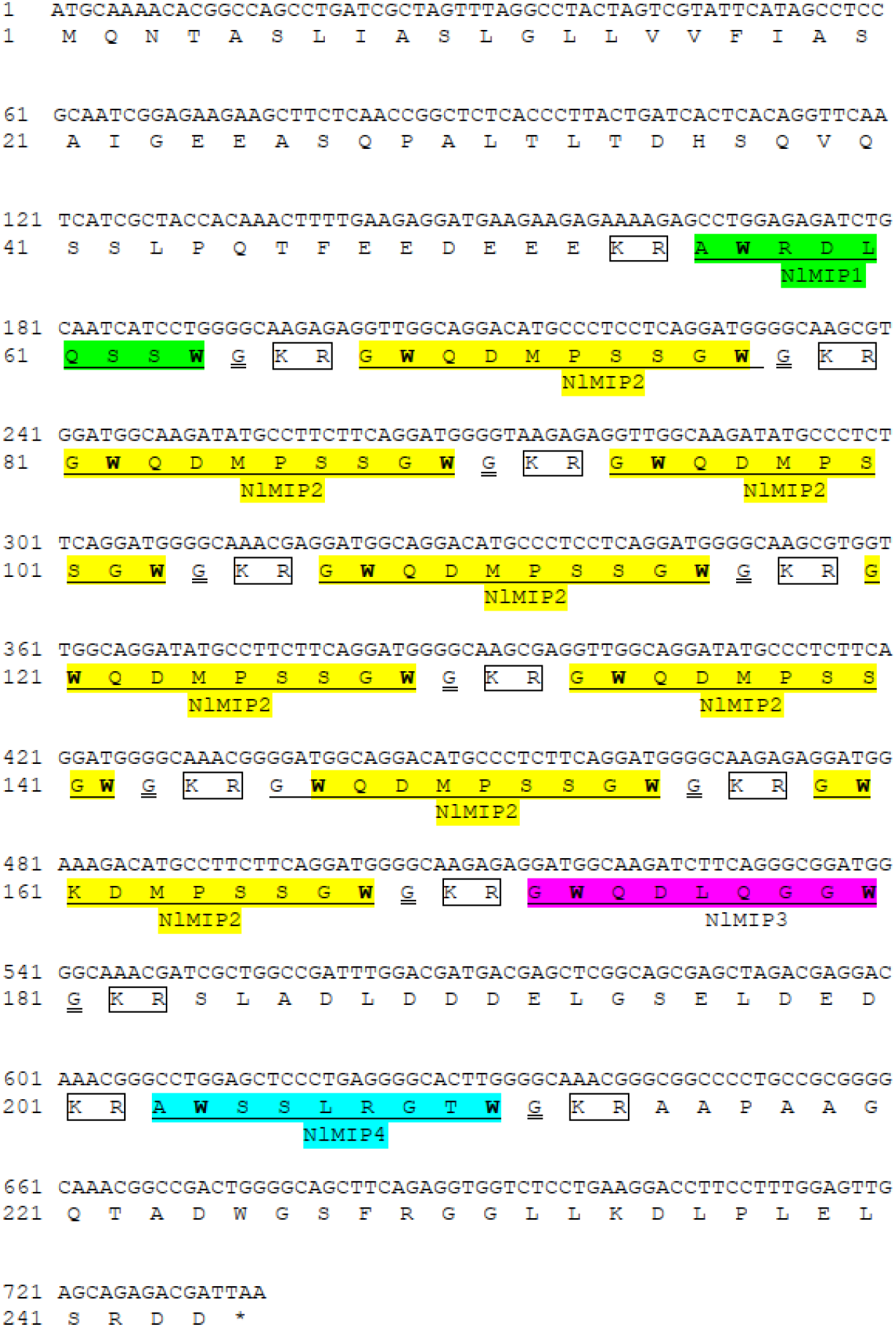
Nucleotide and amino acid sequence of the BPH *mip* precursor gene (*mip)*. The four distinct mature peptides—MIP1 (green), MIP2 (yellow), MIP3 (purple), and MIP4 (cyan) are distinguished by unique colors. Cleavage sites (KR) are denoted by rectangular boxes, while the glycine residues (G) essential for amidation are highlighted by double underlining.

**Figure 5 - figure supplement 2.**
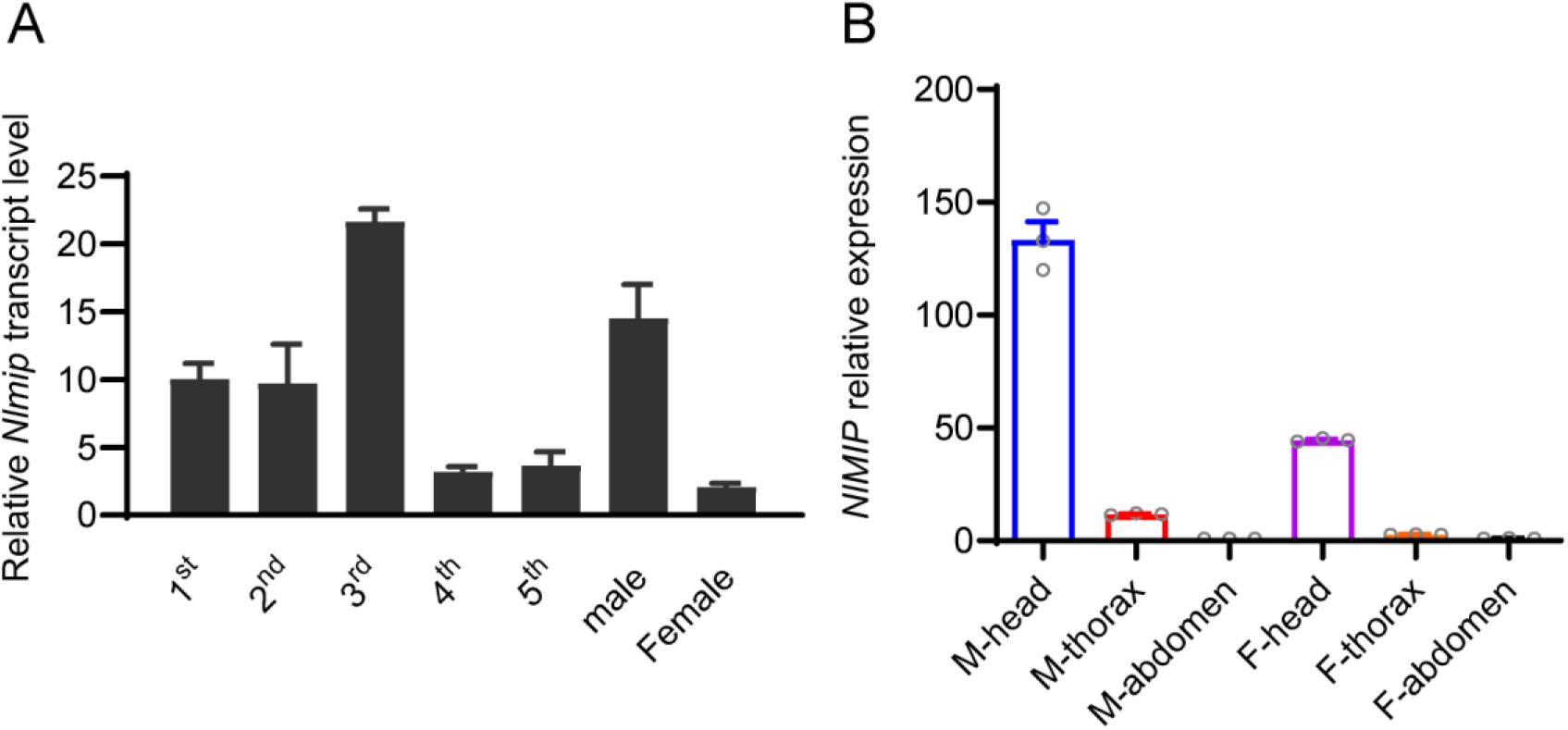
Relative quantification of *mip* transcript levels in different developmental stages (A) and tissues (B) of *N. lugens*. A. Relative expression of *Nlmip* gene in *N. lugens* at different developmental stages. **B.** Relative expression of *Nlmip* gene in *N. lugens* in different tissues. M: male; F: female. Data are shown as mean ± s.e.m.

**Figure 5 - figure supplement 3.**
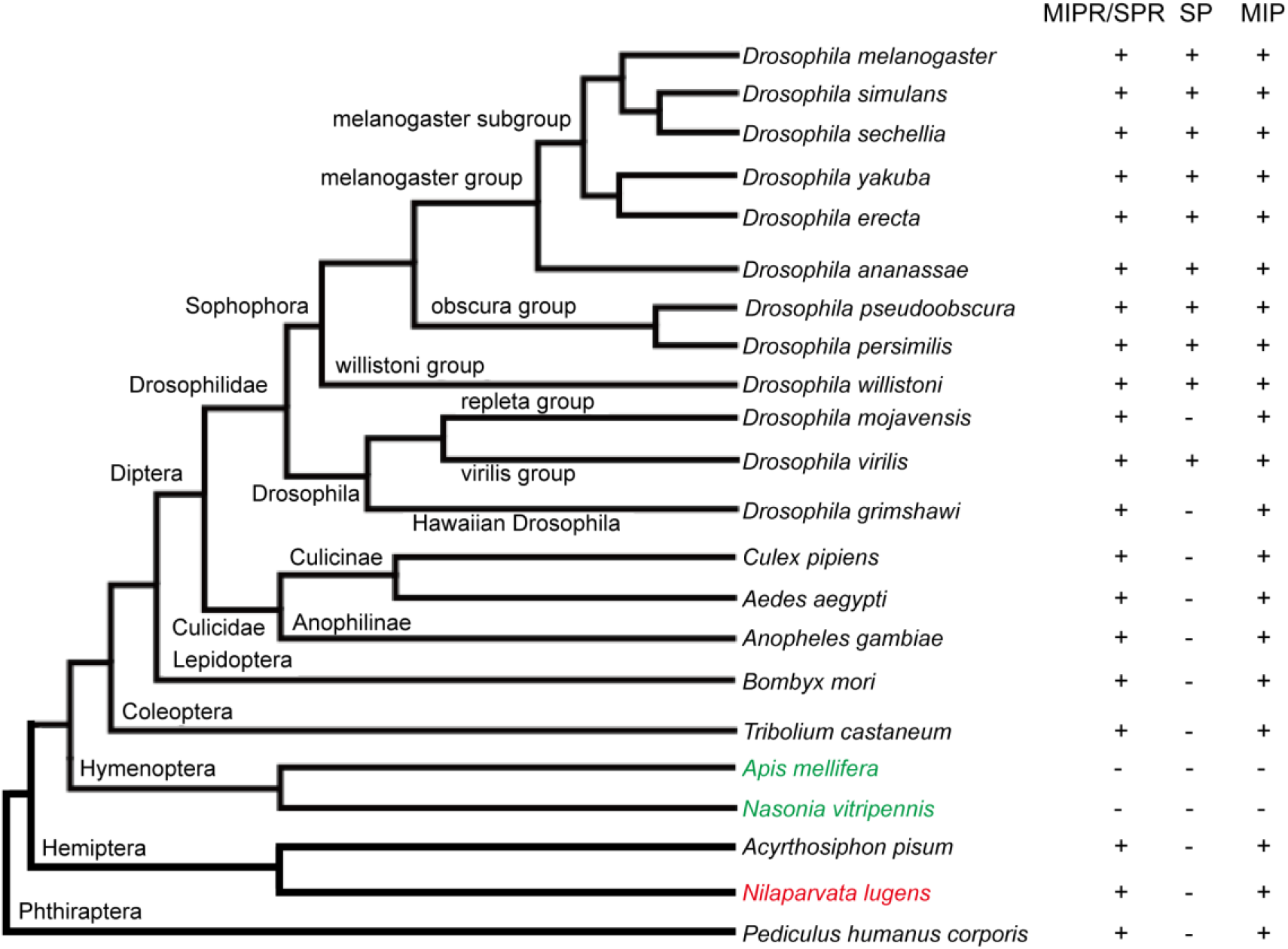
The presence of SPR/MIPR and MIP is widespread across the genomes of most insects that have been sequenced, while the presence of SP is restricted to certain species within the *Drosophila* genus. The brown planthopper (marked in red) lacks SP. In Hymenopteran insects (marked in green), the honey bee *Apis mellifera* and the parasitic wasp *Nasonia vitripennis*, neither SP, MIP, nor SPR/MIPR are found. The "+" symbol represents presence, while the "-" symbol indicates absence. The phylogenetic tree of different insect species has been downloaded and modified from http://flybase.org/blast/.

**Figure 5- figure supplement 4.**
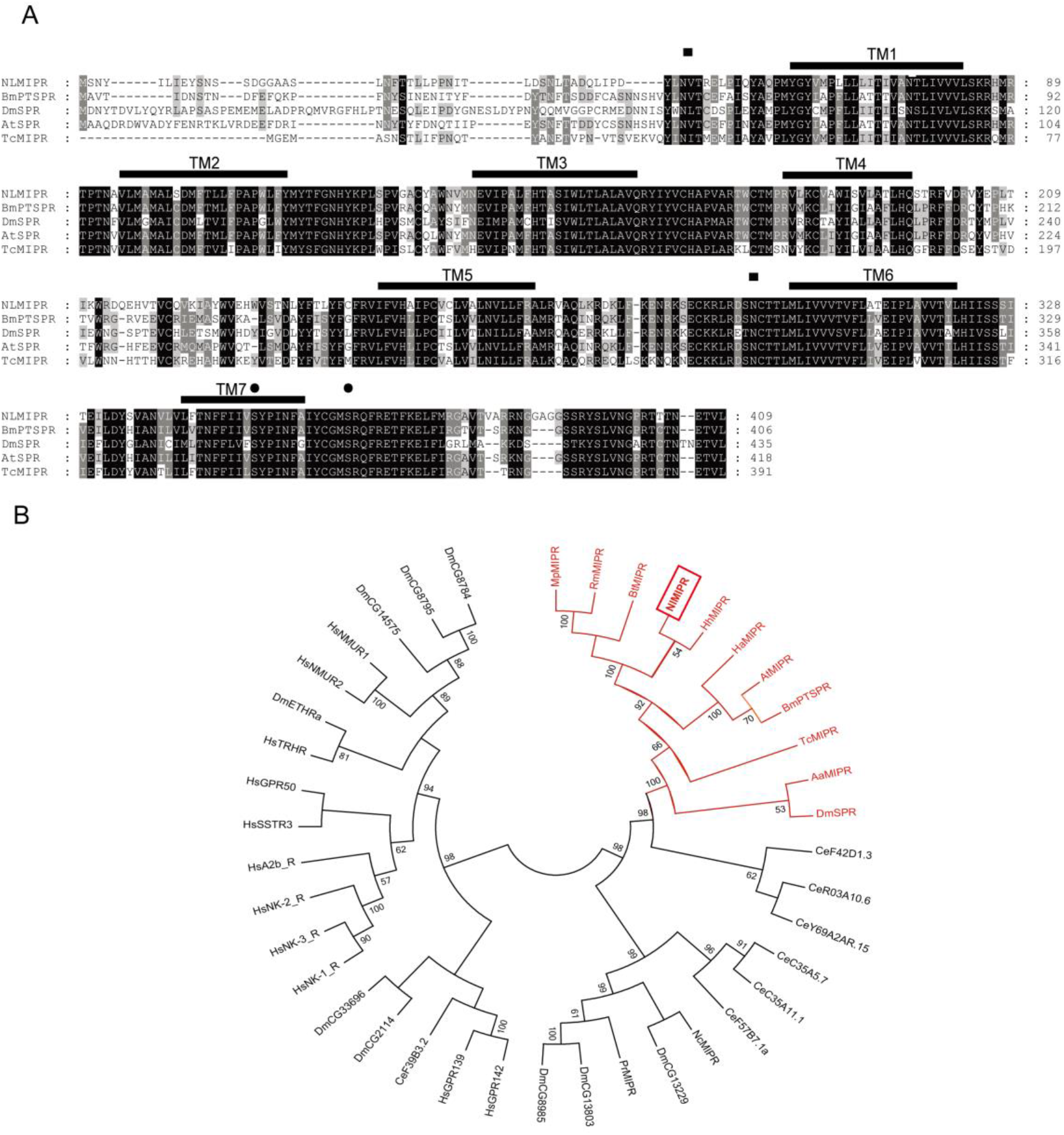
Sequence analysis of MIPRs. **A** Multiple comparison of MIPR in brown planthopper (NLMIPR) and four other insect species (*Bombyx mori*, *Drosophila melanogaster*, *Amyelois transitella* and *Tribolium castaneum*). The black lines (TM1-TM7) depict the transmembrane domains. **B.** Phylogenetic analysis of MIPRs in different insect species.

**Figure 5-figure supplement 5.**
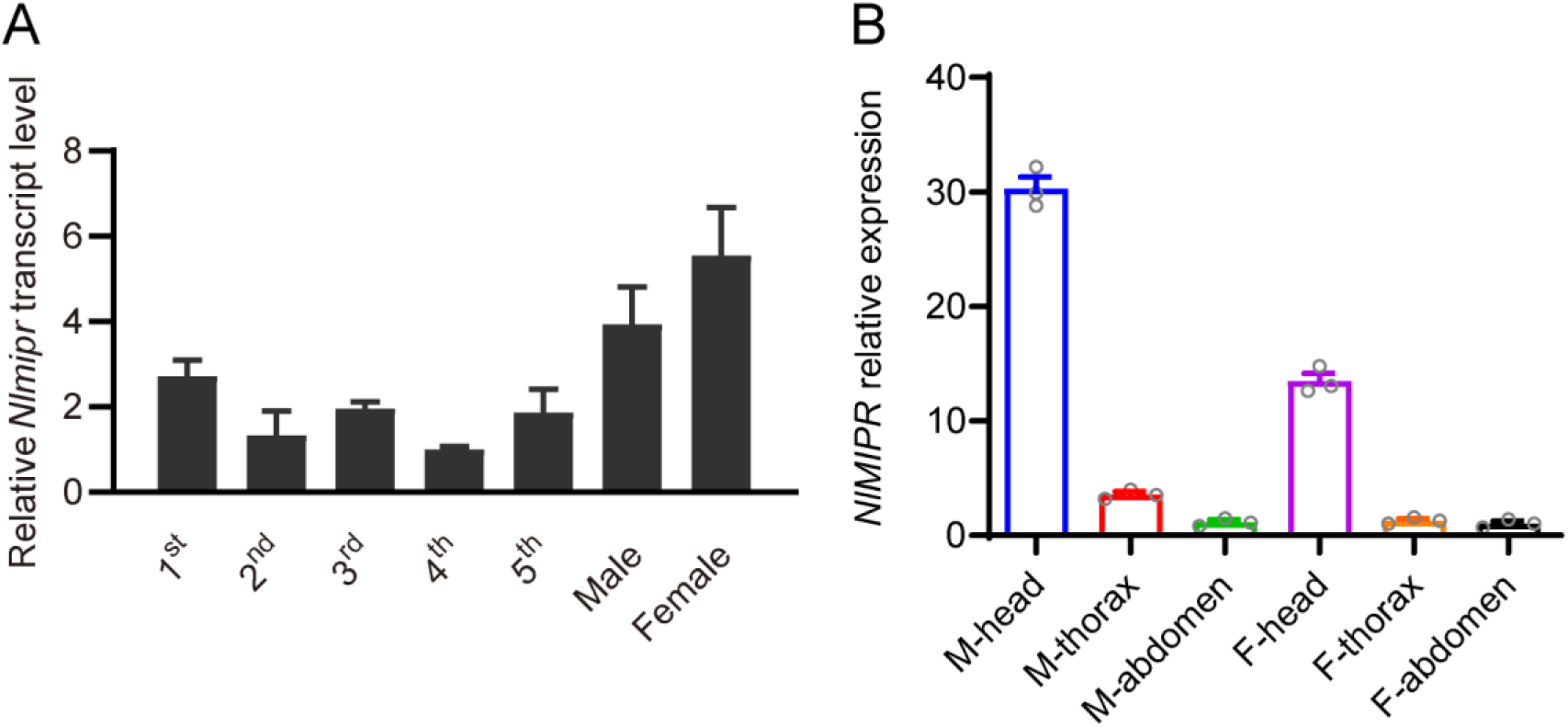
MIP receptor expression in BPH: relative quantification of *mipr* transcript levels in different developmental stages (A) and tissues (B) of *N. lugens*. A. Relative expression of *Nlmipr* gene in *N. lugens* at different developmental stages. **B.** Relative expression of the *Nlmipr* gene in different tissues of adult *N. lugens*. M: male; F: female. Data are shown as mean ± s.e.m.

